# Theory for weakly polydisperse cytoskeleton filaments

**DOI:** 10.1101/2022.01.12.476073

**Authors:** Vadim Warshavsky, Marcelo Marucho

**Affiliations:** Department of Physics and Astronomy, The University of Texas at San Antonio, TX 78249, USA

## Abstract

Cytoskeleton filaments have the extraordinary ability to change conformations dynamically in response to alterations of the number density of actins/tubulin, number density and type of binding agents, and electrolyte concentration. This property is crucial for eukaryotic cells to achieve specific biological functions in different cellular compartments. Conventional approaches on biopolymers solution break down for cytoskeleton filaments because they entail several approximations to treat their polyelectrolyte and mechanical properties. In this article, we introduce a novel density functional theory for polydisperse, semiflexible cytoskeleton filaments. The approach accounts for the equilibrium polymerization kinetics, length and orientation filament distributions, as well as the electrostatic interaction between filaments and the electrolyte. This is essential for cytoskeleton polymerization in different cell compartments generating filaments of different lengths, sometimes long enough to become semiflexible. We characterized the thermodynamics properties of actin filaments in electrolyte aqueous solutions. We calculated the free energy, pressure, chemical potential and second virial coefficient for each filament conformation. We also calculated the phase diagram of actin filaments solution and compared with the corresponding results in in-vitro experiments.

## I. INTRODUCTION

Eukaryotic cells can dynamically regulate the biological environment and the polyelectrolyte and mechanical properties of cytoskeleton filaments to achieve specific biological functions as diverse as directional growth, shape, division, plasticity, and migration [1]. For instance, an increase in the number density of G-actin/tubulin and electrolyte concentration can lead to conformation transformations from orientation disorder (isotropic) to orientation ordered (nematic) phase, as well as increasing the filaments average length. Additionally, a growth in the number density of binding agents, such as divalent ions or linker proteins can yield bundling or network conformations [2]. These self-organization behavior, yet poorly understood, has been observed experimentally. Nowadays, valuable information on the distribution and type of cytoskeleton conformations in cells is obtained from fluorescence and electron microscopy images [3–6]. Whereas, confocal microscopy captures their dynamic conformation changes [7–9]. However, this information usually provides an incomplete molecular understanding of the interplay between polydispersity, semiflexibility, polyelectrolyte, and mechanical properties of cytoskeleton filaments on their conformational dynamics, self-organization, and stability. This understanding is crucial to elucidate the biophysical principles underlying fundamental biological functions of eukaryotic cells in normal and pathological conditions, which may vary depending on the cell type and location, gender, age, and inheritance conditions.

A substantial amount of theoretical research has been done in the field to study the isotropic to nematic phase transformation in macromolecules solution. Conventional understanding of the properties of these polyelectrolytes is based on monodisperse (e.g. same filament lengths), mean-field theories, and rod-like cylindrical filament models (e.g. with contour lengths shorter than their persistence length). These methods break down for cytoskeleton filaments because they entail several approximations to treat the inter-filament interactions, electrolytes, and filament structures. Onsager [10] showed that the expansion of the free energy functional up to the second virial coefficient provides accurate results for monodisperse long charged rods in the coexisting isotropic and nematic phases. In the nematic phase, the free energy is expressed as a functional of angular distribution functions of rods which minimizes the free energy functional. Onsager introduced a trial function for the angular distribution functions depending on a single parameter to increase the efficiency on these calculations. Subsequently, a variety of modifications and extensions of the Onsager’s theory were proposed [11, 12]. For instance, Odijk introduced a Gaussian type trial function for the angular distribution. While less accurate than the Onsager’s approximation, it overcame some limitations on the numerical calculations. It provides explicit, analytic expressions for the thermodynamic properties that indeed facilitated the phase diagram transition and coexistance analysis [13]. A different approach was proposed to estimate the angular distribution functions for monodisperse, long cytoskeleton filament rods [2]. They were not obtained by minimizing the free energy, but postulating a particular form in each phase with a single variational parameter characterizing the width of the angular distributions.

Additionally, corrections were made to the orientational part of Onsager’s free energy to consider monodisperse macromolecules having contour lengths larger or the same order than their persistence length [14, 15]. Furthermore, later approximations were developed to patch the orientational free energies for rigid and semiflexible macromolecules [13, 16]. A different approach was introduced by Sluckin [17] who generalized the Onsager’s theory to describe polydisperse rigid rods having the same Gaussian form for the size distribution function in both isotropic and nematic phases. Moreover, Odijk’s anzatz was generalized to the polydisperse case. where the nematic phase onset was formed by rods with lengths larger than the average length of the size distribution in the isotropic phase. Different modifications of the method were proposed to address this shortcoming [18–21]. In particular, the input size asymmetric distribution functions in the form of Schulz and log-normal distributions were considered.

On the other hand, a particular approach was introduced for amphiphilic micellar suspensions [22]. The size distribution function of micellar macromolecules was not considered as an input value, but rather the standard chemical potential which governed the size-angular distribution functions [23]. It was found that the micella average lengths in the coexisting isotropic and nematic phases are different, with larger micella sizes in the nematic phase.

The size distribution function of uncharged polydisperse actin filament rods in the bundling phase was calculated from a free energy that accounts for hard-core filament repulsion and short-range attractions[24]. It was shown that short-range attractions, arising either from linker proteins, depletion-mediated attractions, or polyvalent ions, enhances the tendency of filaments to align parallel to each other, yielding an increase in the average filament length, and a decrease in the relative width of the distribution of filament lengths.

These approaches for phase diagram studies consider specific macromolecular properties neglecting others. For instance, some approaches focused on polyelectrolyte and polydispersity properties only, whereas, other approximations account for semiflexibility and polyelectrolyte properties, and so on. However, for cytoskeleton filaments, it is imperative to consider the balance and competition between contributions coming from their polydispersity, semiflexibility, polyelectrolyte, and mechanical properties to the total free energy. When accounting for all of these features, one can formulate a more accurate and realistic description of conformational dynamics, self-organization, and stability properties of cytoskeleton filaments in different cell compartments.

As a first step to face this challenge, we introduce in this article a novel density functional theory for polydisperse, semiflexible cytoskeleton filaments. The approach accounts for the equilibrium polymerization kinetics, length and orientation filament distributions, as well as the electrostatic interaction between filaments and the electrolyte. As a unique feature, the formulation is able to determine critical parameter values governing the isotropic-nematic phase diagram behavior. This approach is essential to study the self-organization behavior of actin filaments taking place in different cell compartments, where the G-actin polymerization and electrolyte conditions may generate filaments with different conformations and lengths long enough to become semi-flexible. Specifically, we consider experimental conditions on the actin filaments solution where the isotropic-nematic phase diagram transition is due to changes in the G-actin concentration. Additionally, the filament average size for different G-actin concentrations was fixed by changing the gelsolin proteins concentration [25, 26]. From these special equilibrium conditions, we obtained the size distribution function, whereas we used Sluckin’s trial function to calculate the angular distribution function in the nematic phase. Additionally, we introduced an anzatz for the standard chemical potential excess of actin filaments that results in the asymmetric Schulz’s distribution function for the actin filaments size. This distribution function agrees with those used in light scattering experiments on actin filaments [25]. In addition, we generalized the formula for the orientational free energy introduced in reference [13] to the case of polydisperse system to account for the filament semiflexibility. Finally, we calculated the isotropic-nematic phase diagrams for a variety of Schulz’s size distributions, persistence lengths, and concentrations of monovalent ions in the electrolyte solution. We also compared these results with available experiment data [26].

The paper is organized as follows. The Theory is described in part II, the numerical results are given in part III, the discussion is provided in part IV, and the details of the calculations are presented in Appendix.

## II. MATERIALS AND METHODS

We consider a solution of polydisperse actin filaments, each of them has a length *L* = *l_m_v*, where *l_m_* is the length of an actin monomer unit, and *v* the number of monomer units representing the filament size. Each filament has its own direction in 3D space, which is characterized by a body-angle *ω* accounted from the chosen nematic director. In spherical coordinates, we have *dω* = sin *θdθdφ* with the *z* direction taken along the nematic axis.

The size-angular density distribution function of actin filaments *ρ_ν_*(*ω*) is given by the following expression

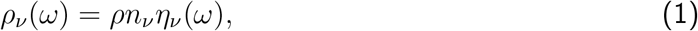

where *ρ* = *N/V* is a total density of filaments, *N* is a total filaments number of any size, and *V* represents the volume of the system. Basically, *ρ_ν_*(*ω*) represents the number of filaments of size *v* oriented along the direction *ω*. Additionally, *n_ν_* is the size distribution function averaged over all angles, and *η_ν_*(*ω*) represents the angular distribution functions, which also depend on the filament size *v*. The normalization conditions for the size- and angular-distribution functions *n_ν_* and *η_ν_*(*ω*) are

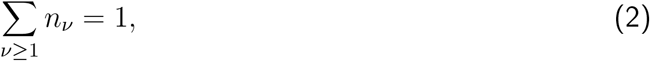

and

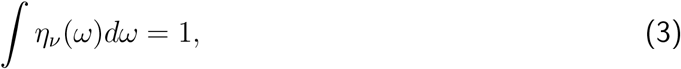

respectively. The summation in Eq.(2) is performed over all filament sizes *v*, whereas the integration in Eq.(3) is carried out over the whole body angle. The size distribution function *n_ν_* is characterized by the filament average size < *v* >, namely the average degree of polymerization, as well as the normalized standard deviation *σ*, such as

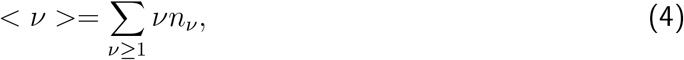

and

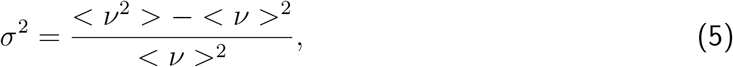

where

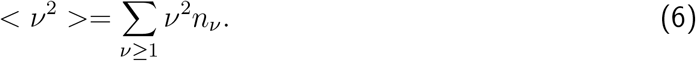

### A. The free energy density functional

We define the dimensionless Helmholtz free energy of cytoskeleton filaments *f* as follow

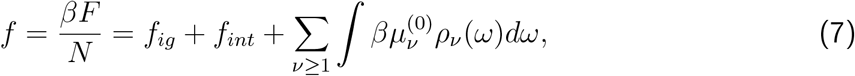

where *F* is the free energy, *β* = 1/*k_B_T* the inverse thermal energy, *k_B_* the Boltzmann constant, *T* the temperature, *f_ig_* the dimensionless ideal gas free energy, *f_int_* the dimensionless energy due to the inter-filament interactions, and 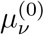 the standard chemical potential of *v*-sized filaments.

The expressions for *f_ig_* and *f_int_* as functionals of the density distributions *n_ν_* and *η_ν_*(*ω*) are provided below.

#### 1. Ideal-gas free energy f_ig_

The ideal-gas free energy per volume of the system *V* can be written in the following form

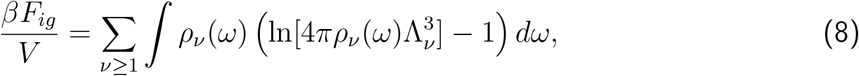

where 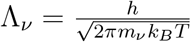 is the thermal deBroglie wavelength of *ν*-sized filaments, *h* is the Planck constant, and *m_ν_* = *νm*_1_ stands for the *ν*-sized filament mass. Substitution of Eq.(1) into Eq.(8) yields

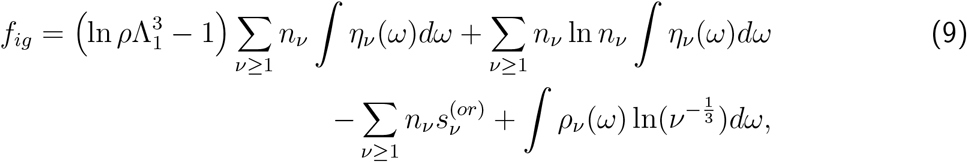

where *f_ig_* = *βF_ig_/N* is the ideal-gas free energy per filament 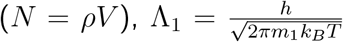 the monomer thermal deBroglie wavelength, and 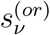 the filament orientational entropy

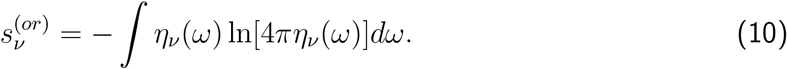

#### 2. Interaction free energy f_int_

We write the interaction free energy per volume *V* in a mean-field fashion, namely

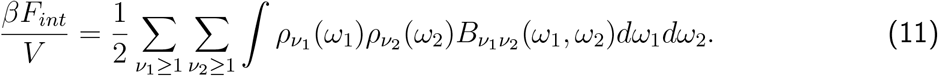

where *B*_*v*_1_*v*_2__(*ω*_1_,*ω*_2_) represents the cluster integral

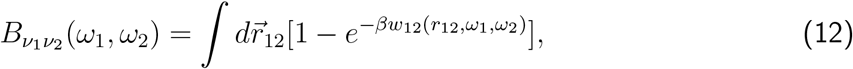

In Eq.(12), *w*_12_ is the interaction potential between two filaments, and *r*_12_ the closest distance between them. The expression (11) is a good approximation for the interaction free energy of monodisperse rigid charged rods. In fact, it becomes exact in the limit of infinitely long rods, i.e. for *L/D* → ∞ [10].

Substitution of Eq.(1) into Eq.(11) yields the following expression for *F_int_* as a functional of the densities *n_ν_* and *η_ν_*(*ω*)

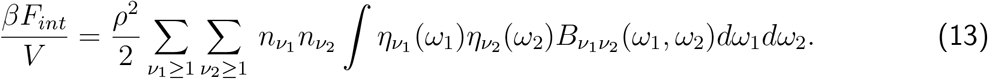

To calculate the cluster integral *B*_*v*_1_*v*_2__(*ω*_1_,*ω*_2_) in obvious form, we model the interaction potential between two charged rod filaments *w*_12_(*r*_12_, *ω*_1_, *ω*_2_) as follow

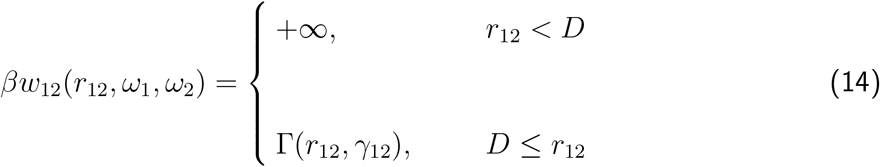

where *D* is the bare rod diameter, and *γ*_12_ the angle between them. The function Γ(*r*_12_, *γ*_12_) represents the following repulsive electrostatic inter-filament interaction potential [27]

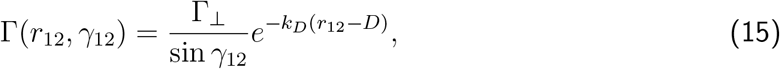

where

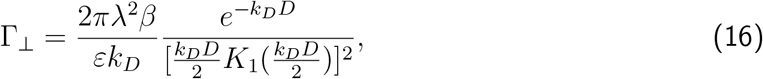

In Eq.(16), 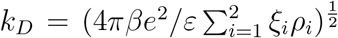 stands for the inverse Debye length, *K*_1_ the modified Bessel function of second kind of first order, λ the filament linear charge density, *ε* the water solvent dielectric permittivity, *e* the electron charge, and *ξ_i_* and *ρ_i_* are the ion valency and concentration of species *i* in solution, respectively. Substitution of Eqs.(14) and (15) into Eq.(12) results in the following expression for the cluster integral *B*_*v*_1_*v*_2__ (*ω*_1_,*ω*_2_)

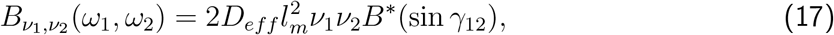

where *D_eff_* and *B*\*(sin *γ*_12_) are the effective diameter and the dimensionless function given by Eqs.(A12) and (A14), respectively. The details on these calculations are presented in Appendix A.

Furthermore, substitution of Eq.(17) into Eq.(13) provides the following form for the interaction free energy

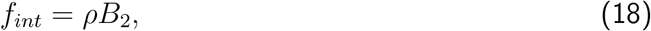

where

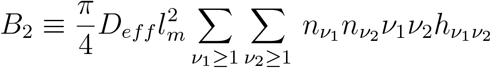

is the extension of the second virial coefficient for polydisperse filaments, and the function *h*_*v*_1_*v*_2__ is given by

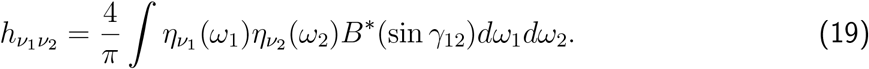

We note that the parameter *h*_*v*_1_*v*_2__ accounts for the orientational averaged contributions coming from the exclude volume and charge density interactions between filaments of size *v*_1_ and *v*_2_, as well as the influence of the ionic strength on the second virial coefficient.

### B. The distribution functions

We introduce the Lagrange functional

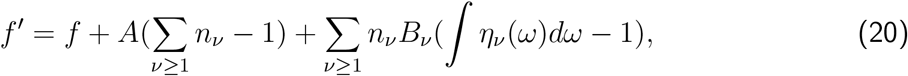

to find the distribution functions, where *f* represents the free energy functional given by Eq.(7), whereas *A* and *B_ν_* are the Lagrange multipliers accounting for the normalization conditions given by Eqs.(2) and (3), respectively. For a fixed angular distribution function *η_ν_*(*ω*), the equilibrium size distribution function *n_ν_* minimizes the Lagrange functional. Thus, we have

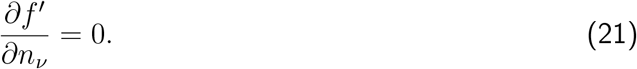

Using the chain rule, the l.h.s of Eq.(21) can be rewritten in the following form

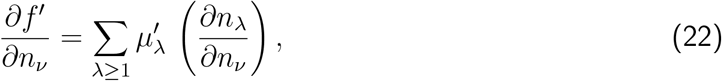

where

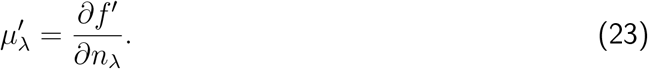

An additional relationship for the distribution functions {*n_ν_*} is coming from Eq.(4). Differentiation on both sides of Eq.(4) yields

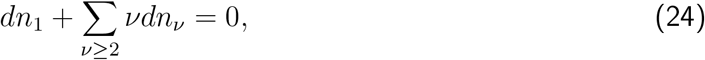

As a result, Eq.(22) can be written as follow

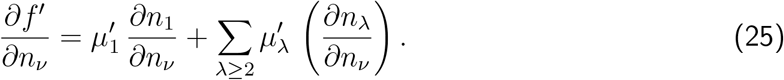

Substitution of Eq.(24) into the r.h.s of Eq.(25) leads to

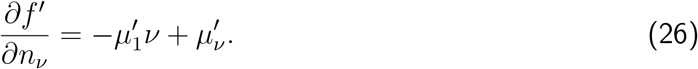

Consequently, Eqs.(21) and (26) yield

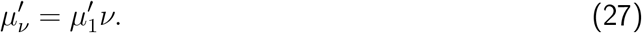

It is worth noting that *v*-sized filaments can be considered as separated ’pseudo-phases’. Additionally, the ’pseudo-phases’ with all possible sizes *ν* are in equilibrium with each other if the equilibrium condition given by Eq.(27) is executed. In fact, the value for 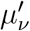 in Eq.(27) does not represent the real chemical potential of *v*-sized filaments since it is coming from the Lagrange functional *f*′ in Eq.(23) rather than the Helmholtz free energy *f*. To calculate 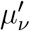, we substitute Eq.(20) into Eq.(23) and use Eqs.(3),(7),(9), and (18). We obtain

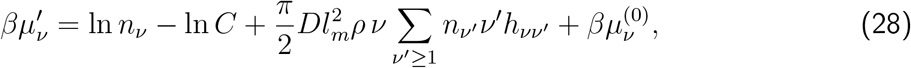

where *C* is the following constant

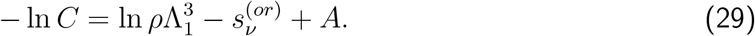

We substitute the expression (28) into the condition of equilibrium (27) to get the following equation for the length distribution function *n_ν_*

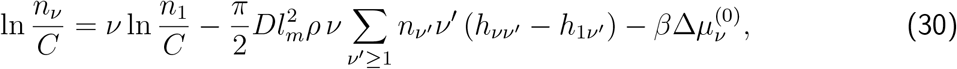

where 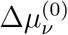 denotes the standard chemical potential difference between *ν* actin units aggregated in a *v*-mer filament 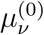 and the one in the single dispersed phase 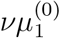, i.e.

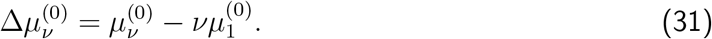

Finally, we substitute the parameter 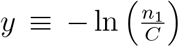 into Eq.(30) to get the following master equation for *n_ν_*

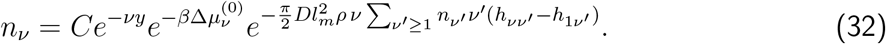

Similarly, for a fixed distribution function *n_ν_*, the angular distribution function *η_ν_*(*ω*) is obtained by using the variational principle

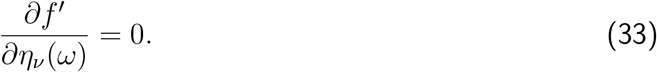

Substitution of Eq.(20) into Eq.(33) and the use of Eqs.(2),(4),(7),(9),(10),(18), and (19) yield

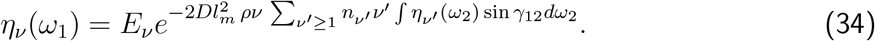

where *E_ν_* is a constant that satisfies the following relationship

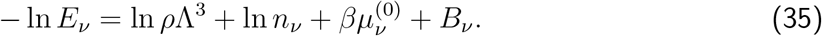

To obtain the expression for *E_ν_* we integrate both sides of Eq.(34) with respect to *ω*_1_ and use the normalization conditions (2) and (3). Finally, we replace the expression obtained for *E_ν_* into Eq.(34) to get the following master equation for *η_ν_*(*ω*)

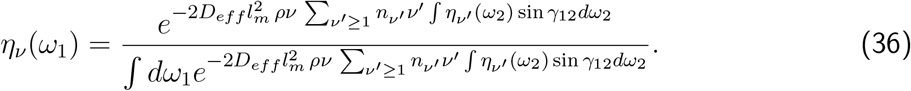

Furthermore, the angular distribution function *η_ν_*(*ω*) depends only on the polar angle *θ* due to the symmetry of the system. Thus, we have *η_ν_*(*ω*) = *η_ν_*(*θ*).

To find the size and angular distribution functions *n_ν_* and *η_ν_*(*θ*), we solve the expressions (32) and (36) using the successive iterations method. To this end, we choose the monodisperse size distribution as the initial guess for the first iteration step, i.e. *n_ν_* = *δ*(*v* – < *v* >), where *δ* is a Dirac delta-function. Substitution of this expression for *n_ν_* into Eq.(36) generates the following equation to numerically calculate *η*(*θ*) [28]

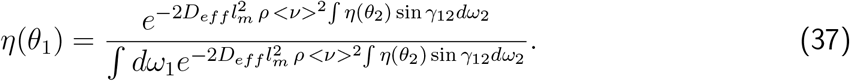

More efficient but less accurate values for the monodisperse angular distribution function *η*(*θ*) were proposed using some functional forms. For instance, Onsager [10] introduced the trial function *η*(*θ*) = *α* cosh(*α* cos *θ*)/(4π sinh *α*). Another commonly used although slightly less accurate anzatz is the so-called Odijk’s trial function [13]

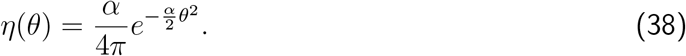

We note that these trial functions depend on a single unknown parameter *α* which is chosen to minimize the free energy functional *f*.

In the second iterative step, we substitute the angular distribution function *η*(*θ*) obtained in the first step into Eq.(19) to calculate *n_ν_*. Since *η*(*θ*) does not depend on *ν*, it follows from Eq.(19) that *h_νν′_* – *h*_1*ν*′_ = 0. Additionally, we use in Eq.(32) the anzatz

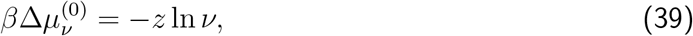

to obtain a size distribution function *n_ν_* in the asymmetric Schulz-Zimm form [29]

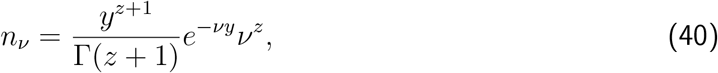

where *z* represents the conventional polydispersity parameter, whereas 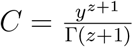 is the constant coming from the normalization condition given by Eq.(2). In appendix B, we show that the polydispersity parameter *z* is related to the normalized standard deviation of the size distribution function 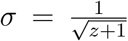, whereas the parameter *y* depends on the values < *ν* > and *z* only, i.e *y* =(*z* + 1)/ < *ν* >.

Finally, substitution of Eq.(40) into Eq.(36) gives an equation to calculate the function *η_ν_*(*θ*). Getting the numerical solution of this equation indeed requires a high computational cost because it depends on two variables *v* and *θ*. Alternatively, we generalize the monodisperse Odijk’s trial function given by Eq.(38) to the polydisperse case. Specifically, we introduce the following parametrization for the polydisperse angular distribution function [17]

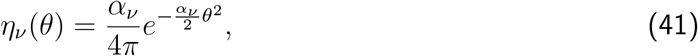

where the polydisperse Gaussian parameter *α_ν_* depends on the filament size *ν*. For weak polydispersity of the system, the ratio 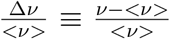 can be considered a small parameter. As a result, the polydisperse Gaussian parameter *α_ν_* can be calculated using the following linear expansion around the monodisperse solution

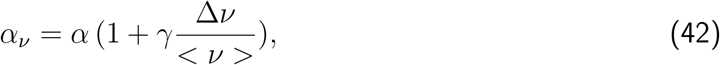

where the two unknown parameters *α* and *γ* are chosen to minimize the free energy functional *f*. In the monodisperse limit, we have that Δ*v* → 0 and *α_ν_* → *α*.

### C. Free energy in the nematic phase

We substitute the normalization conditions (2) and (3) into Eq.(9) to write the Helmholtz free energy per filament as follow

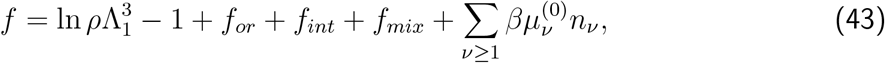

where 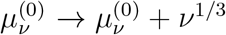, and the terms *f_or_*, *f_int_*, *f_mix_* represent the orientational, interaction, and mixing free energies, respectively. The expressions for these energies are provided below.

#### 1. Orientational free energy

The orientational free energy in Eq.(43) is given by the expression

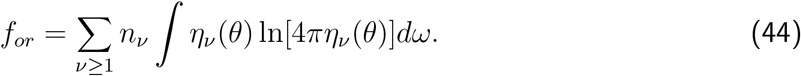

Substitution of the expressions for *η_ν_*(*θ*) (41) and (42) into Eq.(44) yields

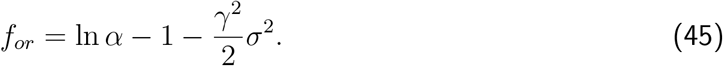

Eq.(45) represents the generalization of the orientational free energy expression obtained by Odijk for monodisperse rods, i.e. for filaments with contour length < *L* > << *P*, where *P* is the filament persistence length. Indeed, in the monodisperse limit *σ* → 0 the correct limit *f_or_* → ln *α* – 1 is obtained for any value of the parameter *γ*[13]. For flexible filaments, where < *L* > >> *P*, the orientational free energy can be written in the following form [13]

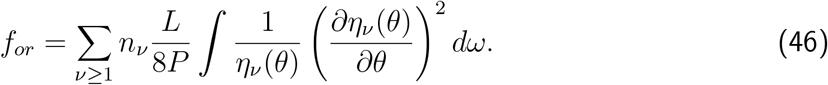

Substitution of the expressions for *η_ν_*(*θ*) (41) and (42) into Eq.(46) leads to

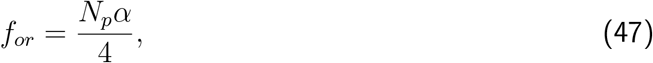

where

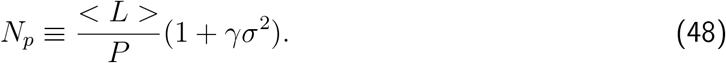

and < *L* >= *l_m_* < *v* >. Certainly, in the monodisperse limit, *σ* → 0 the correct limit 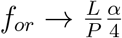 is obtained[11].

To calculate the orientational free energy for any persistence length, we combine these two asymptotic cases for *f_or_* using the following interpolating formula

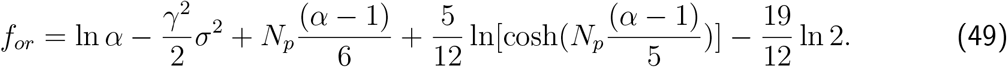

The details on these calculations are provided in Appendix C.

#### 2. Interaction free energy

We substitute Eqs.(40), (41), and (42) into Eqs.(18) and (19) to get

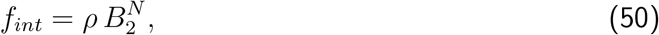

where

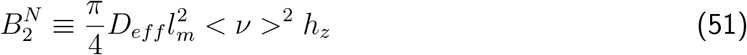

represents the second virial coefficient for polydisperse filaments in the nematic phase, and

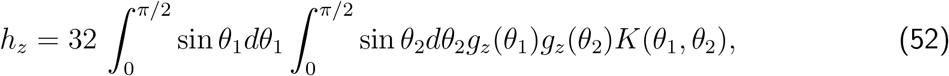

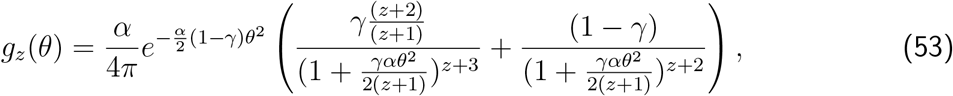

and

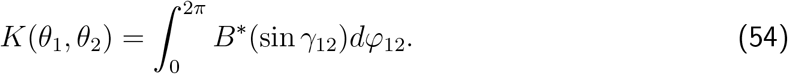

The details on these calculations are presented in Appendix D.

Eq.(50) represents the generalization of the interaction free energy expression for polydisperse actin filaments. In fact, in the monodisperse limit *z* → ∞ the expression for *g_z_*(*θ*) goes to

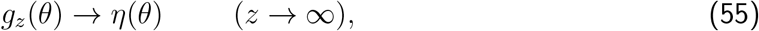

which is the monodisperse distribution function given by Eq.(38). Similarly, Eqs.(50), (52), (53), (54) recover the correct expression for the filaments interaction free energy *f_int_* in the monodisperse case

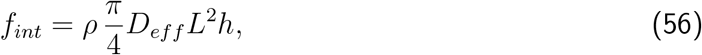

with

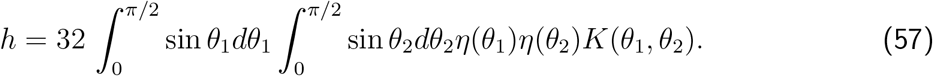

#### 3. Mixing free energy

The mixing free energy in Eq.(43) can be written as

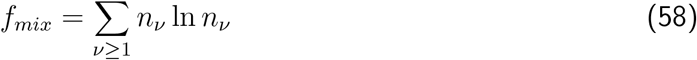

This expression is not well defined in the monodisperse limit (*z* → ∞) since the density distribution *n_ν_* → *δ*(*v* – < *v* >) and the mixing free energy diverges rather than going to zero. To overcome this shortcoming, we extract the density *ρ* independent term *f_mix_*(*z* → ∞) in Eq.(E7) from the r.h.s of Eq.(58). The resulting renormalized mixing free energy reads

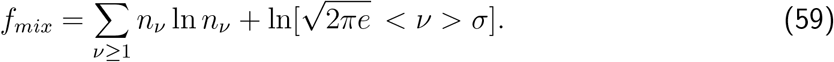

This new expression (59) recovers the correct value in the monodisperse limit, i.e. *f_mix_* → 0 for *σ* → 0 (*z* → ∞). More details on these calculations are provided in Appendix E.

### D. Free energy in the isotropic phase

In the isotropic phase, there is no preferential direction for the filament orientations, thus, the orientational distribution function *η_ν_*(*θ*) = 1/(4π) for any filament length, and the orientational free energy becomes

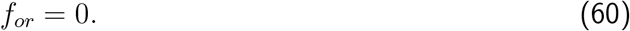

The expression for the interaction free energy is obtained from Eqs.(18),(19) and (4). We have

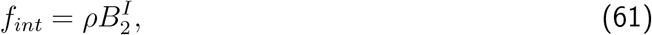

where 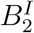 is the second virial coefficient for polydisperse filaments in the isotropic phase

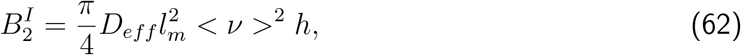

with

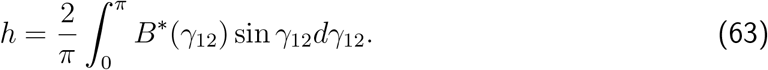

Finally, substitution of Eq.(A14) into Eq.(63) generates the following expression for *h* in isotropic phase

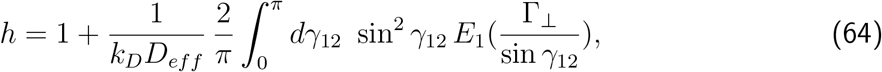

where *E*_1_(*x*) is the elliptic integral given by Eq.(A9).

### E. Isotropic-nematic phase at equilibrium

The isotropic-nematic phase equilibrium in a system of polydisperse macromolecules occurs when the chemical potentials of each species *μ_ν_* and the osmotic pressures Π are equal each others. Specifically, we have [17, 19, 20]

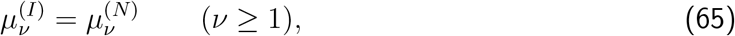

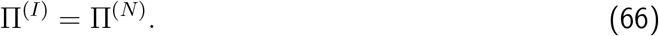

In the case of self-aggregation of actin filaments, there is an additional condition for the chemical equilibrium in each thermodynamic phase, namely

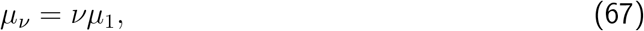

where 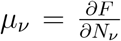 is the chemical potential of *v*-sized filaments, and *μ*_1_ the chemical potential of actin monomers. The condition (67) is obtained using a method similar to the one leading to Eq.(27). Substitution of Eq.(67) into Eq.(65) provides a single equilibrium condition for the chemical potentials, i.e.

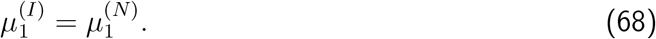

To calculate the chemical potential of actin monomers *μ*_1_ we write the Gibbs free energy of mixture of actin filaments of all sizes in the following form

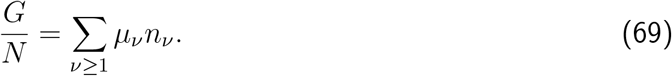

Substitution of Eqs.(4) and (67), and the use of condition 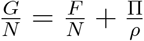 into Eq.(69) yields

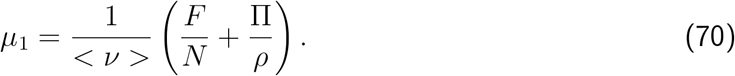

In Eqs.(66) and (70), we use the formula 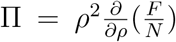 for practical calculations of the osmotic pressure.

Using Eq.(4), we can also write the average filaments length in Eq.(70) as follows

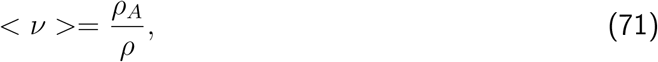

where 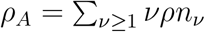 is the G-actin number density. As a result, the average filaments length represents the ratio of *G*-actin and *F*-actin concentrations. Additionally, *G*-actin is usually polymerized in in-vitro experiments in the presence of gelsolin, an actin-binding protein to be known to cap and sever actin filaments [25, 26, 30]. It was noticed in these experiments that the concentration of *F*-actin in the solution becomes almost equal to the concentration of gelsolin, i.e. *ρ* ≈ *ρ_G_*. As a result, the average length of filaments (average degree of polymerization) < *v* > in these experiments was regulated by the gelsolin concentration. In Ref.[26] the variation of concentration of *G*-actin *ρ_A_* was accompanied by a change in the concentration of gelsolin *ρ_G_* to keep the average length of filaments < *v* > fixed during the isotropic-nematic phase transition. Using this experimental condition in our theory, the mixing free energy *f_mix_* and the term with 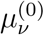 in rhs of expression for free energy (43) are also fixed values and, thus, do not contribute to the phase coexistence properties.

Thus, using Eqs.(43), (50) and (61), the dimensionless free energy *f* in the nematic and isotropic phases can be written as follow

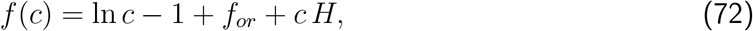

where *c* represents the dimensionless filaments density

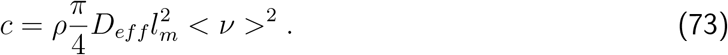

The expressions to calculate *f_or_* are given by Eqs.(49) and (60) for the nematic and isotropic phases, respectively. Whereas, the corresponding expressions for *H* are given by Eqs (52) and (64).

In the nematic phase, the free energy *f* (72) is a function of the parameters of the angular distribution function *α*, *γ* and the dimensionless filaments density *c*, thus, *f* ≡ *f*(*c*; *α, γ*). By minimizing *f*, we obtain the equilibrium parameter functions *α_eq_*(*c*) and *γ_eq_*(*c*) for each value of *c*. Thus, the expression for the nematic free energy at equilibrium becomes *f_eq_*(*c*) = *f* (*c, α_eq_*(*c*), *γ_eq_*(*c*)). The minimization is carried out using the downhill simplex method in multi-dimensions [31]. Finally, the coexisting dimensionless densities *c_I_* and *c_N_* are obtained from the coexistence conditions (66) and (68), whereas the corresponding actin densities 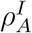 and 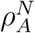 are obtained from Eqs.(71) and (73).

## III. RESULTS

In this section we apply the approach to investigate the isotropic-nematic phase diagram for polydisperse actin filaments in monovalent salt solutions. We considered typical experimental values for key parameters, such as the filament persistence length, the average filament size, the polydispersity parameter, and the electrolyte concentration to elucidate their impact on the orientational and size distributions. We also calculated the thermodynamic properties of the system, including the free energy, pressure, and chemical potential, and consequently, the range of G-actin concentrations and filaments average lengths leading to conformation transformations from orientation disorder (isotropic) to orientation ordered (nematic) phase. In the numerical calculations we chose the values for the filaments diameter *D* = 80*Å*, the monomer units length *l_m_* = 27*Å*, the actin molar weight *m_A_* = 42*kDa*, and the linear filament charge density 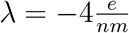.

We calculated the isotropic and nematic coexisting dimensionless densities for rigid monodisperse rods with length *L* = 1 *μm* for the electrolyte’s concentration *c_e_* = 0.1 *M* to compare our results with those obtained by Borukhov et al [2]. Our result is *c_I_* = 3.82 and *c_N_* = 5.10,whereas the corresponding one in Ref.[2] is *c_I_* = 4.25 and *c_N_* = 9.98 (see Fig.3 in Ref.[2]). The source of such discrepancy is associated with the cone approximation used in Ref.[2] for the angular distribution function.

### A. Distribution functions

In this section we present the analysis on the distribution functions *n_ν_* and *η_ν_*(*θ*) in the I-N phase coexistence for polydisperse, semiflexible filaments. We used a persistence length *P* = 18 *μm* and an electrolyte’s concentration *c_e_* = 0.1 *M*. In Fig.1 we plotted the Schulz’s length distribution function *n_ν_* for different average sizes < *ν* > in a weakly polydisperse system with normalized standard deviation *σ* = 0.5 (*z* = 3).

**Figure 1.**
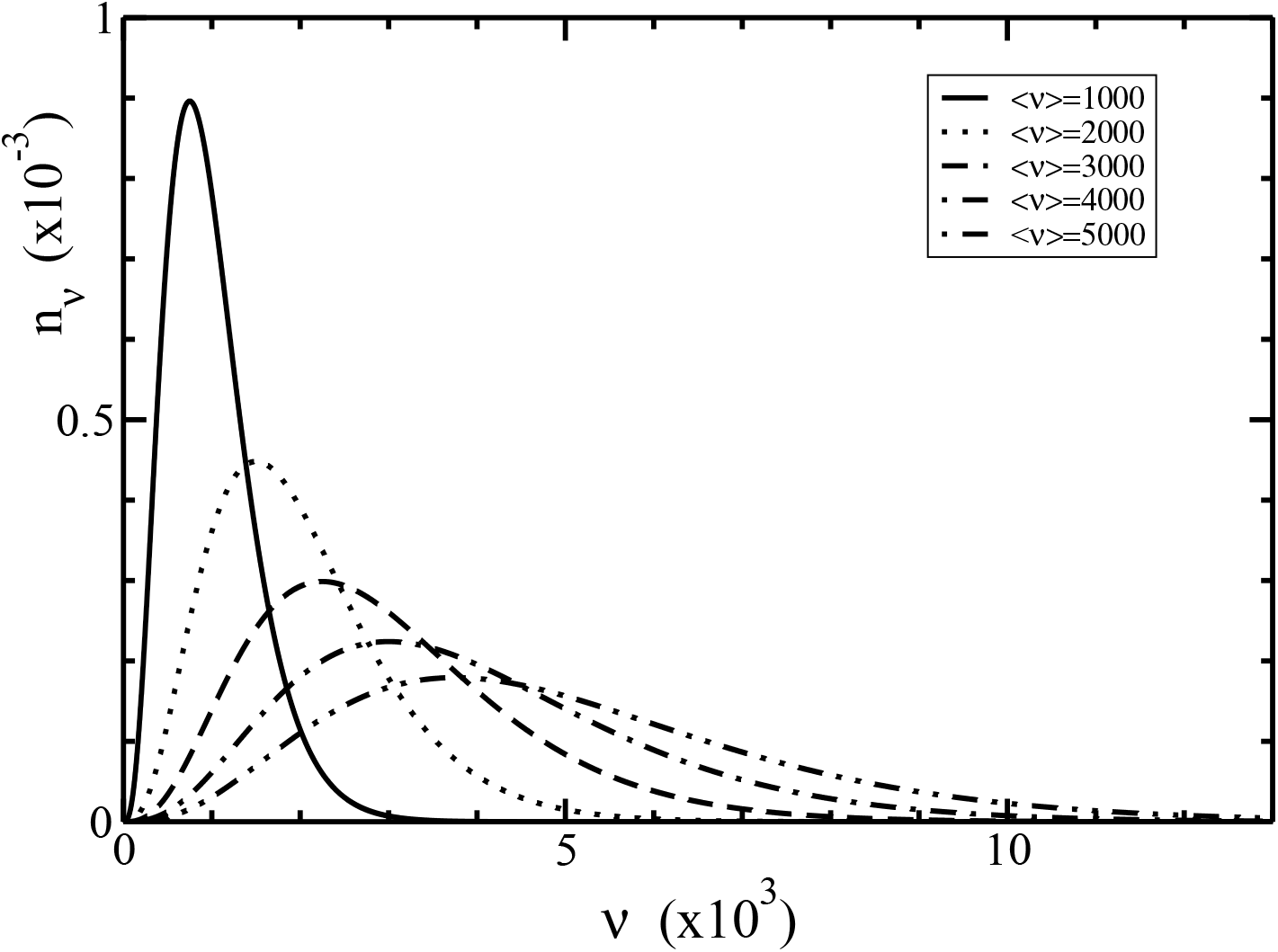
Schulz’s length distribution function *n_ν_* as a function of the filament size *ν* for different average sizes < *ν* > from Eq.(40). The normalized standard deviation is *σ* = 0.5 (*z* = 3).

Our results reveal polydisperse distribution functions with lower peaks and broaden distributions for larger values of the average size < *ν* >. While, the asymmetric behavior characterizes the different increasing rate lengths of barbed and pointed ends. This asymmetry property plays a crucial role on the phase diagram behavior. Since the normalization integral 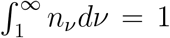 can be split into the sum of two integrals 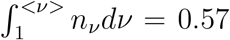 and 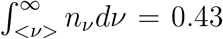 (*z* = 3), each of them is independent of the value < *ν* >, the amount of the filaments with the sizes *ν* shorter than < *ν* > is larger than the one with the sizes longer than < *ν* >. In contrast, monodipserse, symmetric distribution functions are represented by delta-function distributions *n_ν_* → *δ*(*v* – < *ν* >).

In Figures 2 and 3 we depicted the I-N phase coexistence values of the parameters *α* and *γ* appearing in the angular distribution function for different values of the average length < *ν* >.

**Figure 2.**
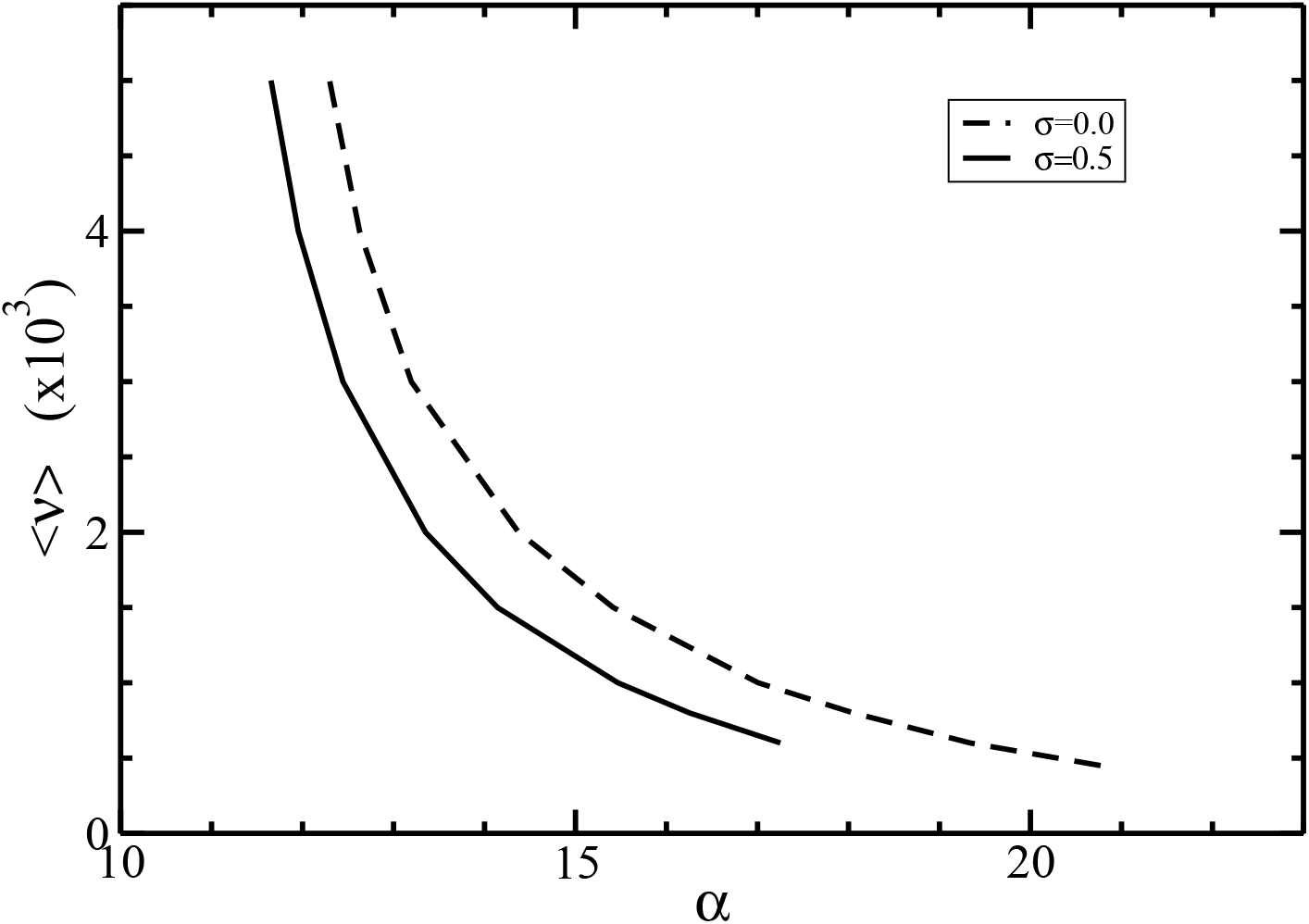
The parameter *α* as a function of the average size < *v* > at the I-N phase coexistence. The normalized standard deviations are *σ* = 0 (dashed line) and 0.5 (solid line).

**Figure 3.**
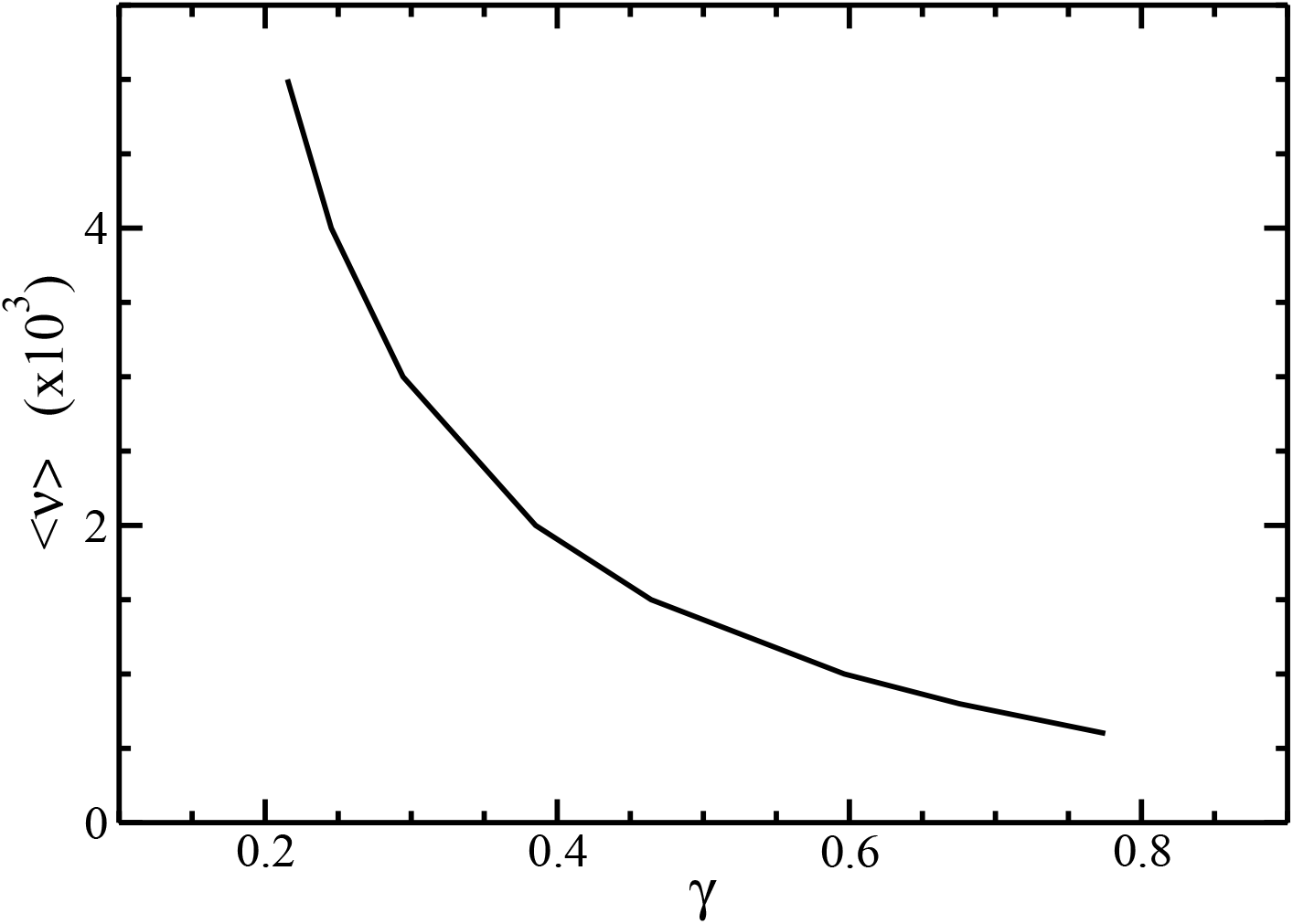
The parameter *γ* as a function of the average size < *ν* > at the I-N phase equilibrium. The normalized standard deviation is *σ* = 0.5.

It is seen that larger average length < *ν* > leads to lower parameters values *α* and *γ*. In fact, larger filament sizes generate narrower angular distributions, increasing the order of the system. For a given value of < *ν* >, Fig.2 also shows that the parameter *α* decreases with increasing the polydispersity parameter *σ*. This is due to the typical broaden size distributions that characterize polydisperse systems, which include short filaments lowering the order of the system.

In Fig.4 we plotted the Gaussian parameter *α_ν_* given in Eq.(42) as a function of the average filament size *ν*. For a fixed size *ν*, the parameter *α_ν_* decreases with increasing the average length < *ν* >. Additionally, the dependence of the orientational distribution function *η_ν_*(*θ*) on the angle *θ* for different sizes *ν* is plotted in Fig.5.

**Figure 4.**
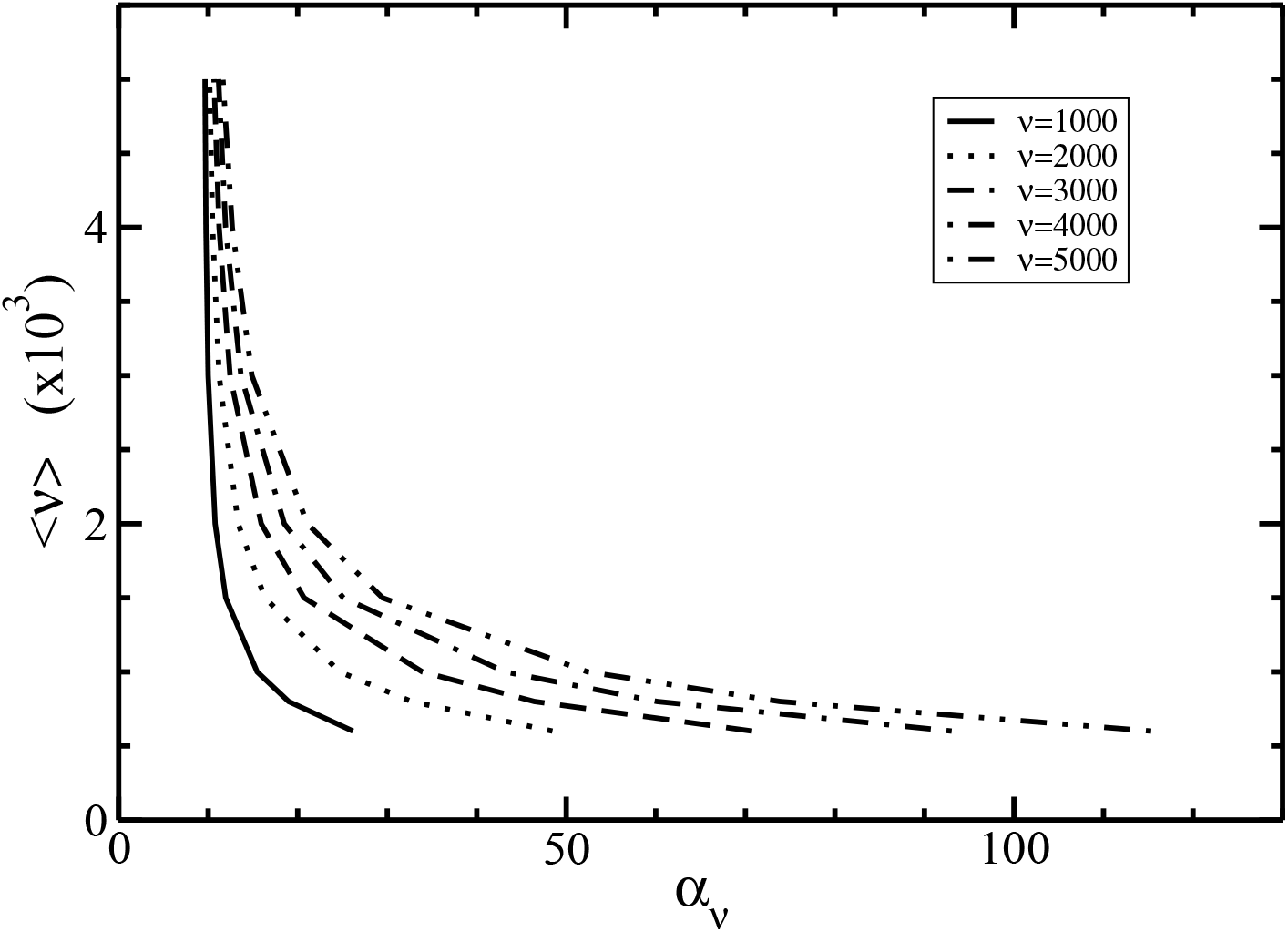
The Gaussian parameter *α_ν_* vs the average size < *ν* > at I-N phase coexistence for different length of filaments *ν*.

**Figure 5.**
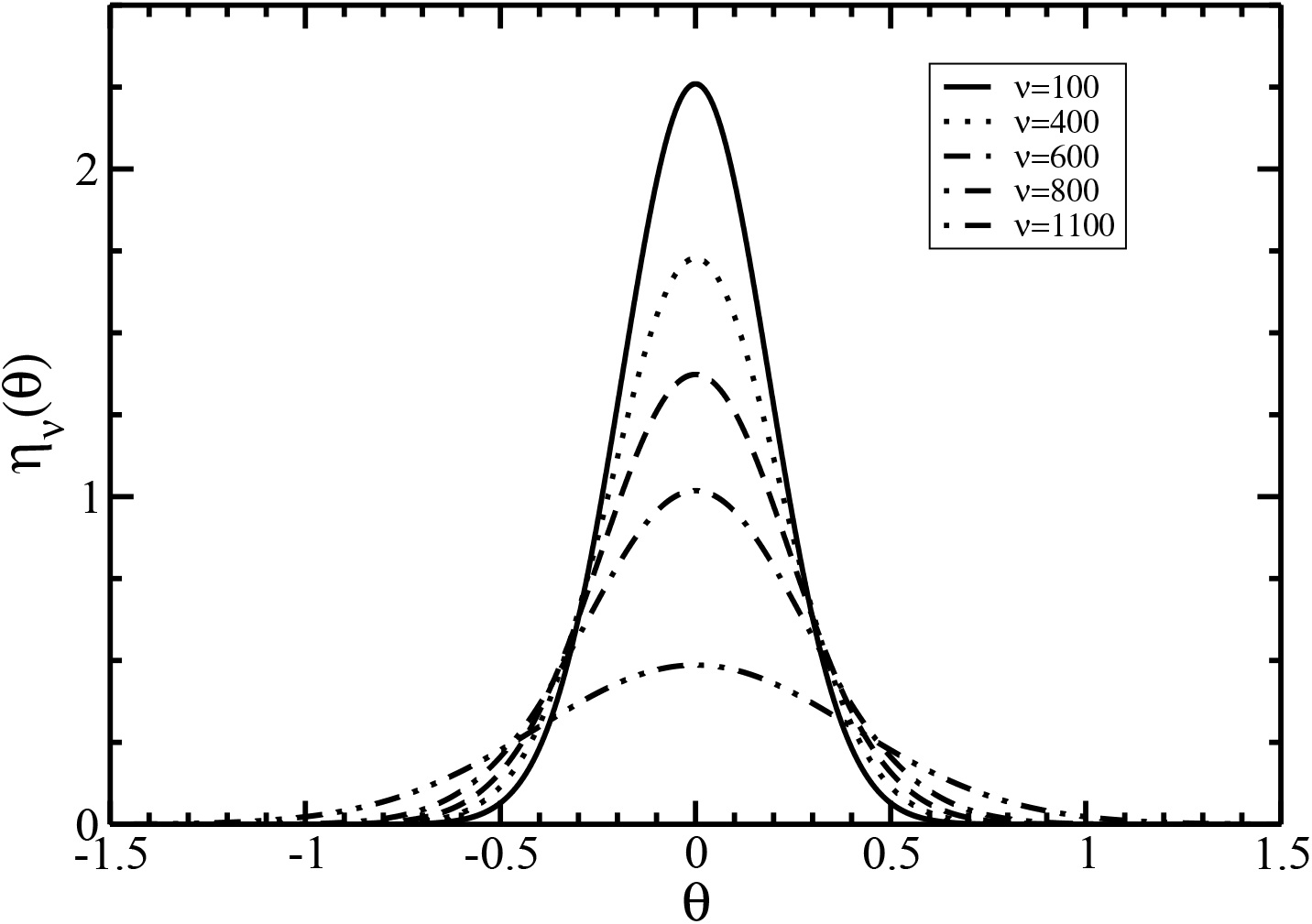
The angular distribution function *η_ν_*(*θ*) as a function of the angle *θ* at the I-N phase coexistence for different sizes *ν*. The average length is < *ν* >= 600.

It is seen that the increase of size *ν* flattens and widens the distribution *η_ν_*(*θ*). On the other hand, the dependence of the total size-angular distribution function *ρ_ν_*(*θ*)/*ρ* = *n_ν_η_ν_*(*θ*) on the filament size *ν* for several angles *θ* is displayed on Fig.6.

**Figure 6.**
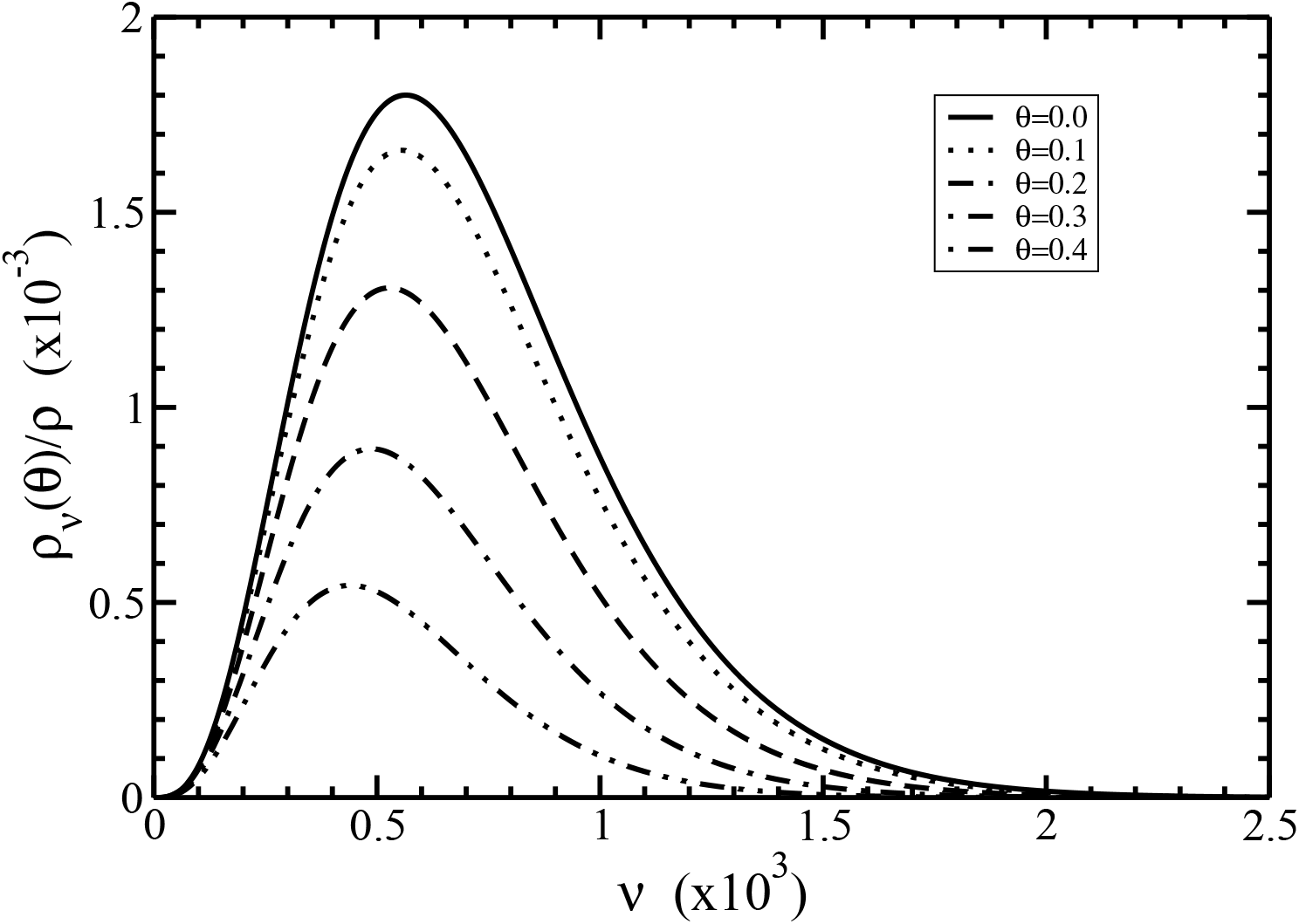
The size-angular distribution function *ρ_ν_*(*θ*)/*ρ* = *n_ν_η_ν_*(*θ*) as a function of the size *ν* at the I-N phase coexistence for different angles *θ*. The average length is < *ν* >= 600.

Our results show that the increase of the angle *θ* flattens the total distribution *ρ_ν_*(*θ*)/*ρ*. This is because most filaments in nematic phase are oriented along the nematic director with *θ* = 0. By increasing the angle *θ*, the amount of filaments decreases, and are directed along the direction with angle *θ*.

### B. Isotropic-nematic phase diagram

In Figure 7 we plotted the I-N phase diagram of actin filaments in term of the coexisting dimensional densities of actin *ρ_A_* for different average lengths < *ν* > and two values for the normalized standard deviation of the Schulz’s distribution function: *σ* = 0 (monodisperse), and *σ* = 0.5 (weakly polydisperse). In these calculations we used the values *P* = 18 *μm* and *c_e_* = 0.1 *M*. In the same figure, we plotted the experimental data extracted from Figures 4 and 5 in Ref.[26].

**Figure 7.**
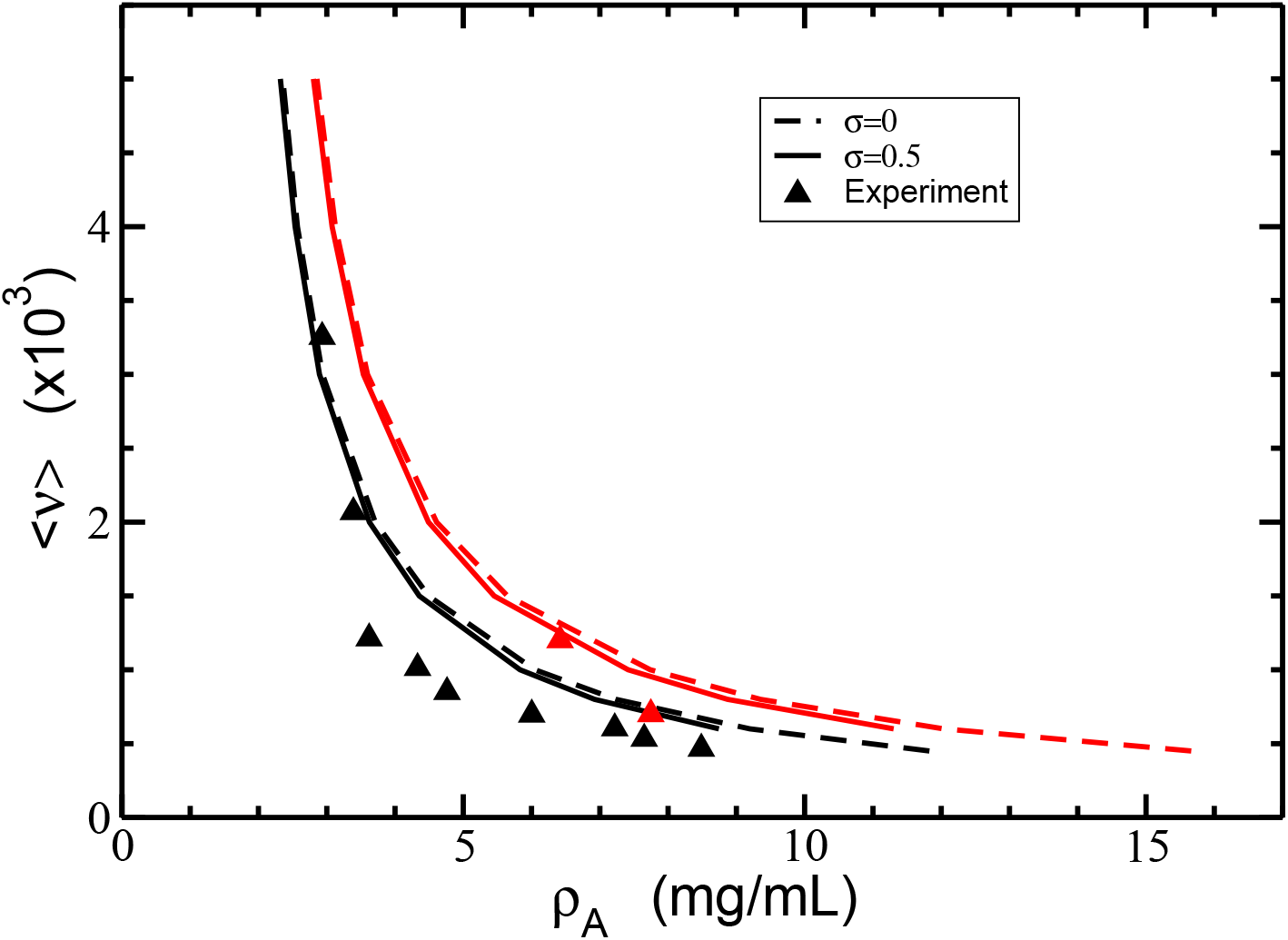
The isotropic-nematic phase diagram in term of coexisting densities of actin *ρ_A_* for different average length of filaments < *ν* > and two values of the normalized standard deviations *σ* = 0 (dashed lines) and *σ* = 0.5 (solid lines). The persistence length is *P* = 18 *μm*, and the electrolyte concentration *c_e_* = 0.1 *M*. The experimental data (triangles) are retrieved from Ref.[26]. The coexisting isotropic and nematic densities are plotted in black and red colors, respectively.

Our results show isotropic and nematic methastable phases represented by the area enclosed between the red and black curves. Whereas, the area to the left of the black curves and to the right of the red curves defines the exclusive randomly oriented (isotropic) and parallel oriented (nematic) phases, respectively. Additionally, the red and black curves display an increase in the average filament length with decreasing G-actin concentration, and consequently, the order of the system. For a given G-actin concentration, the nematic phase generates larger average filament lengths as compared to the isotropic phase. Whereas, higher G-actin concentrations are required in the nematic phase to generate the same average filament length obtained in the isotropic phase. Figure 7 also shows a minor dependence of the coexisting densities on the polydispersity parameter. This is in part due to the experimental condition fixing the same degree of polymerization < *ν* > for both the isotropic and nematic phases. In fact, Fig.7 shows that polydispersity with larger amount of short filaments slightly reduces both isotropic and nematic coexisting densities as compared to the monodisperse case. Thus, additional factors like electrostatic effects compete with hard-body interactions to reduce the coexisting densities in the polydisperse case.

We also studied the polydispersity effects on the nematic (anisotropic) order parameter *S*, which is defined as follow

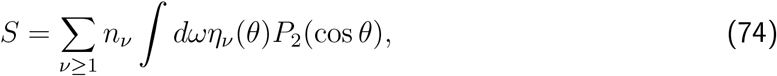

where *P*_2_ = (3 cos^2^ *θ* – 1)/2 is the Legendre polynomial of second order. In the isotropic phase, where *η_ν_*(*θ*) is constant, the order parameter *S* vanishes. In contrary, in a system highly aligned, *η_ν_*(*θ*) becomes the Dirac-delta function leading to *S* = 1. The I-N phase coexistence values of the nematic order parameter *S* for different average sizes < *ν* > and two values of the polydispersity parameter *σ* are displayed in Fig.8. For both values of *σ*, our results reveal a decrease of the order parameter *S* with increasing the average length < *ν* >. Moreover, the order parameter *S* is even smaller for polydisperse filaments.

**Figure 8.**
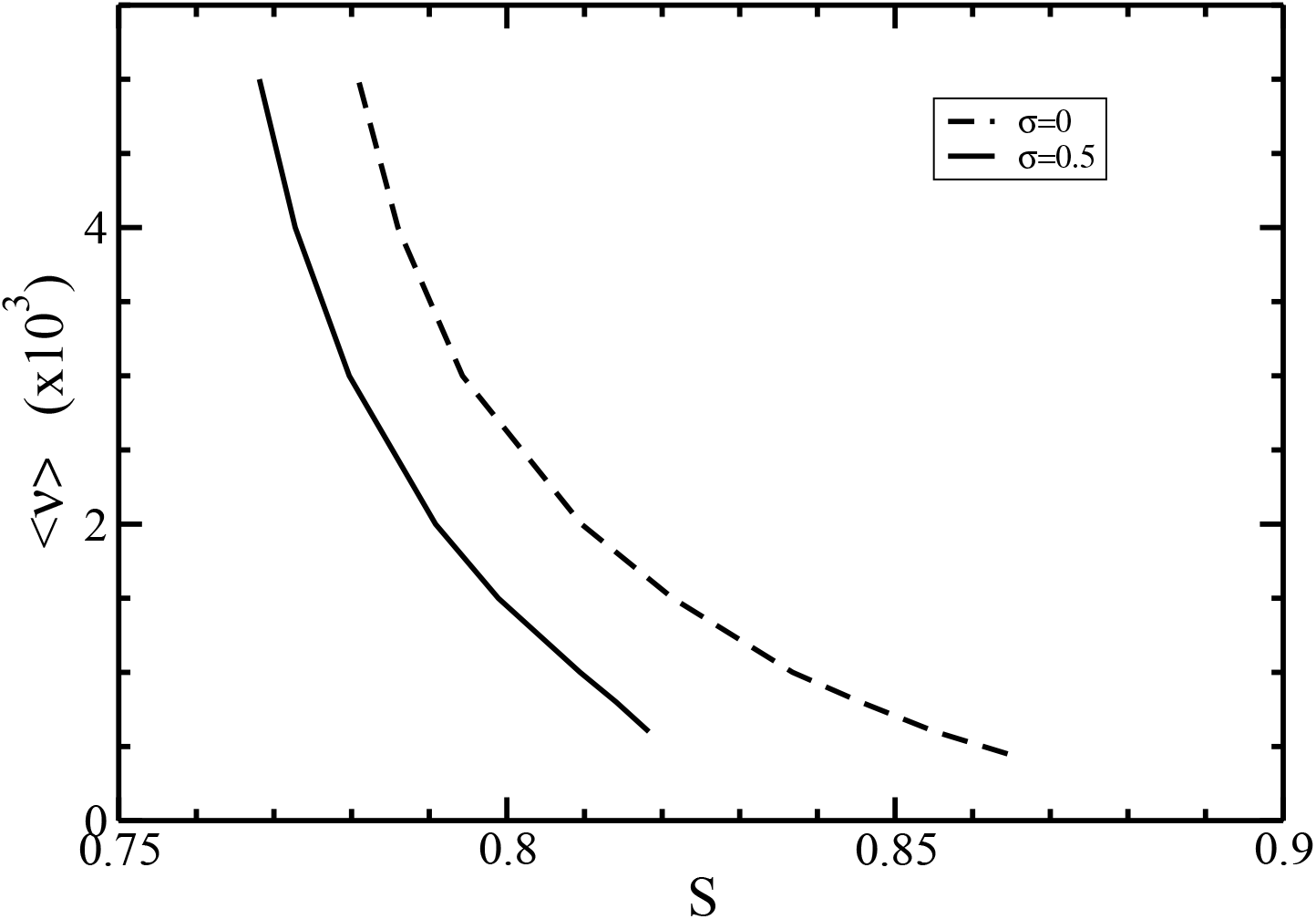
The nematic order parameter *S* at the I-N phase coexistence for different average lengths < *ν* >. The normalized standard deviation are *σ* = 0 (dashed line) and 0.5 (solid line).

Overall, we realized that the orientational order of filaments at I-N phase equilibrium is due to the competition between two contributions. In the previous section, we showed that the increase of average length < *ν* > leads to narrower angular distributions, and thus, higher order in the filaments orientation. On the other hand, Fig.7 displays a decrease in the coexisting nematic density lowering their order. Since the order parameter *S* accounts for both contributions, and the total order becomes smaller for larger average length < *ν* >, we conclude that the contribution coming from the decrease in the coexisting nematic density dominates the order behavior of the filaments over those arising from narrowed angular distributions.

We also considered several filament semiflexibility parameter values since they vary depending on the polymerization buffers, experimental protocols and techniques. We analyzed in Fig.9 the isotropic-nematic phase diagram for two typical values of the persistence length, namely *P* = 7*μm* and *P* = 18*μm*, in physiological conditions (electrolyte concentration *c_e_* = 0.1*M*).

**Figure 9.**
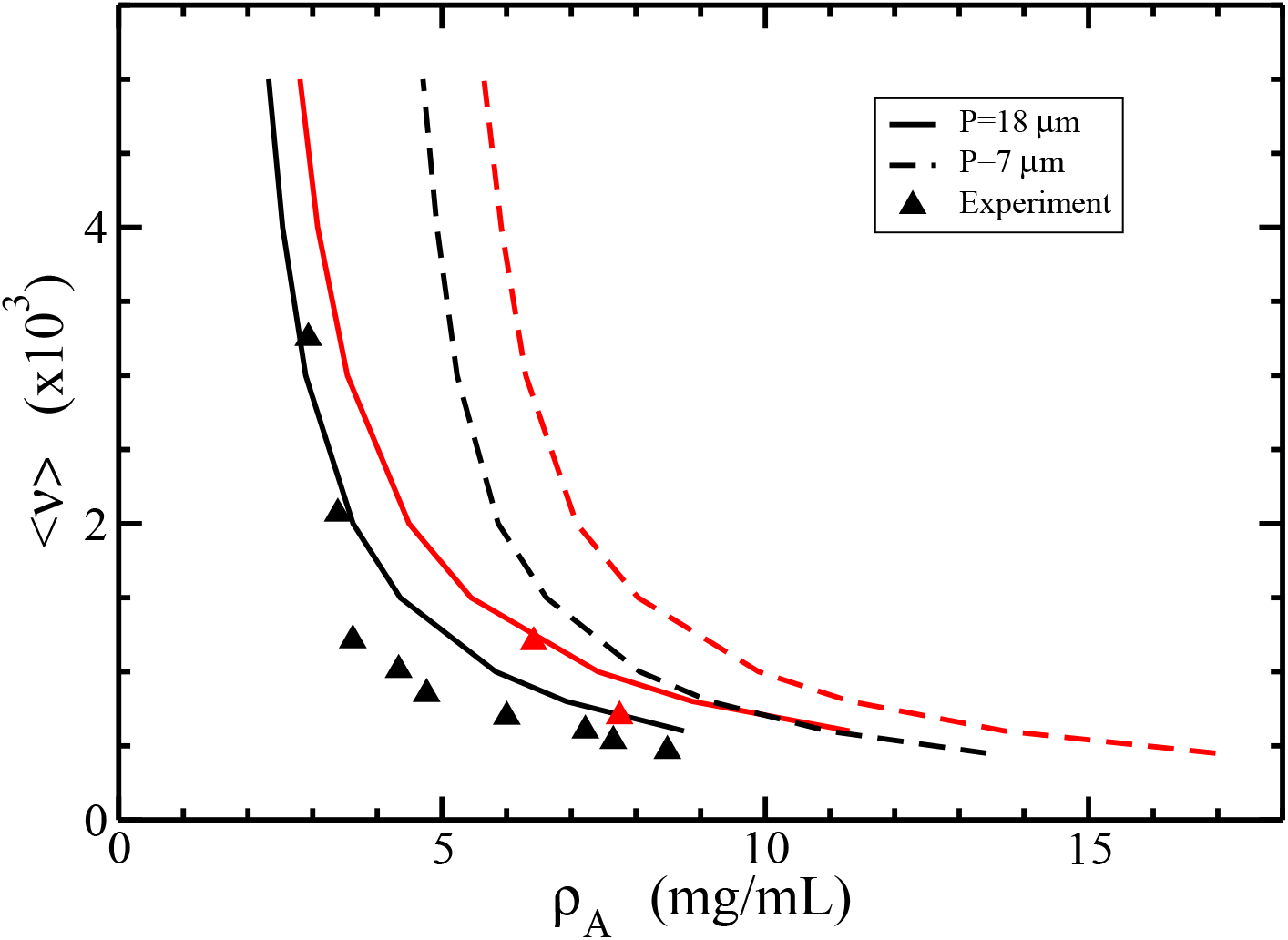
The isotropic-nematic phase diagram in term of coexisting densities of actin *ρ_A_* for different average length of filaments < *ν* > and two values of persistence lengths *P* = 7*μm* (dashed lines) and *P* = 18*μm* (solid lines). The normalized standard deviation is *σ* = 0.5, and the electrolyte concentration *c_e_* = 0.1*M*. The experimental data (triangles) are retrieved from Ref.[26]. The coexisting isotropic and nematic densities are plotted in black and red colors, respectively.

For a given value < *ν* >, we found that larger values of the persistence length *P* decrease both the isotropic and nematic coexisting densities. Additionally, the results for *P* = 18*μm* gives the best fit against the experimental data. This result stems from the impact of the semiflexibility on the orientational free energy *f_or_*. It can be seen from Eq.(49) that the increase of the persistence length *P* leads to the decrease of the orientational term *f_or_*, which boosts the formation of the nematic state and generates smaller values for the coexisting densities.

Finally we studied the effects of the electrolyte’s concentration on I-N phase diagram of actin filaments. We plotted in Fig.10 the dependence of the effective diameter *D_eff_* on the concentration of monovalent ions *c_e_*.

**Figure 10.**
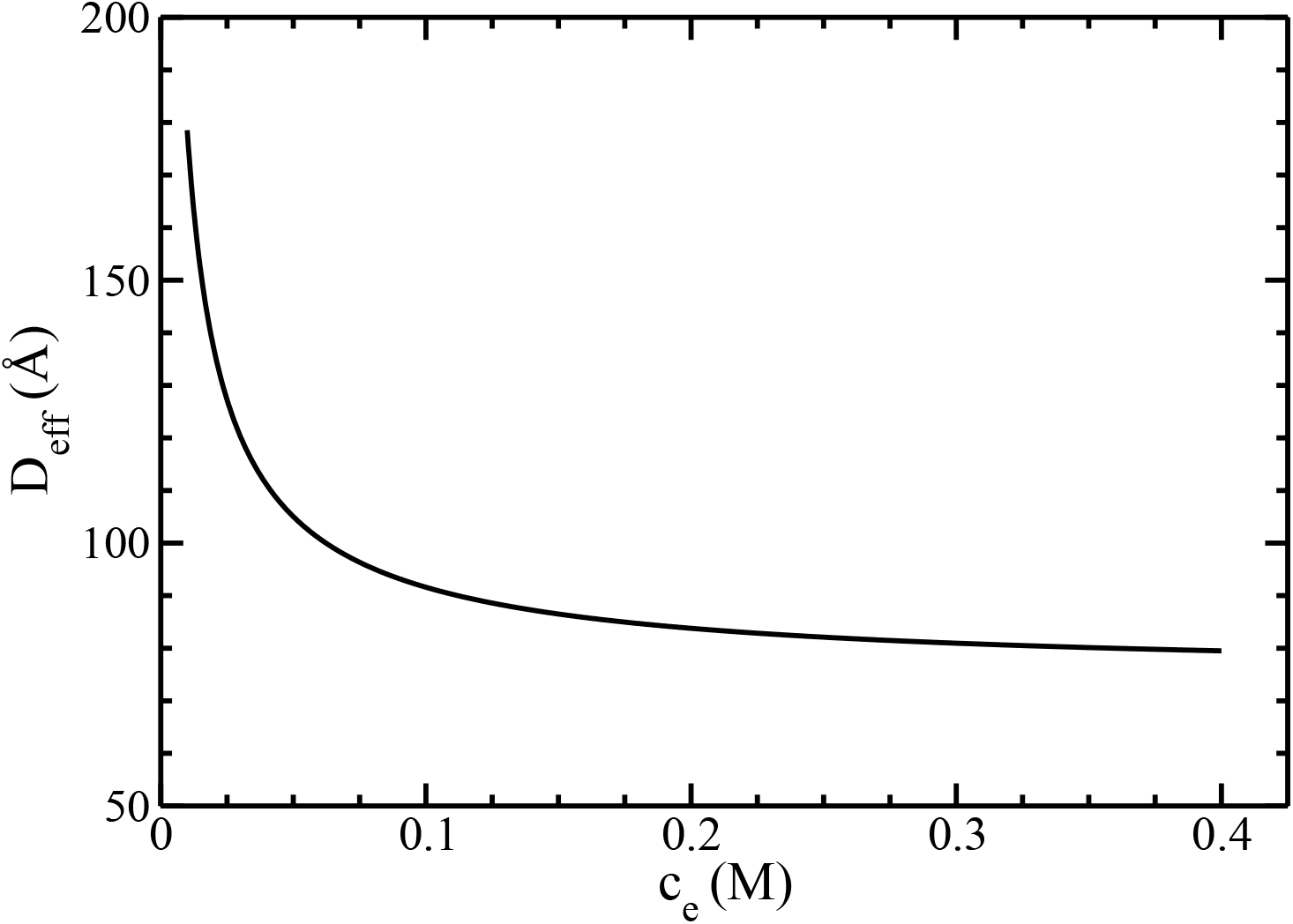
The effective diameter *D_eff_* as a function of the monovalent ions concentration *c_e_*.

Our results show an exponential decay of *D_eff_* with increasing *c_e_*. This result agrees with the formation of an electrical double layer accumulating many counterions around the filament surface. The lower the electrolyte concentration, the larger its thickness and, consequently, the effective filament diameter.

Furthermore, we show in Fig.11 the isotropic-nematic phase diagram for two values of electrolyte concentration *c_e_* = 0.01*M* and *c_e_* = 0.1*M*. It is seen that the increase in electrolyte concentration leads to the increase in both coexisting isotropic and nematic densities.

**Figure 11.**
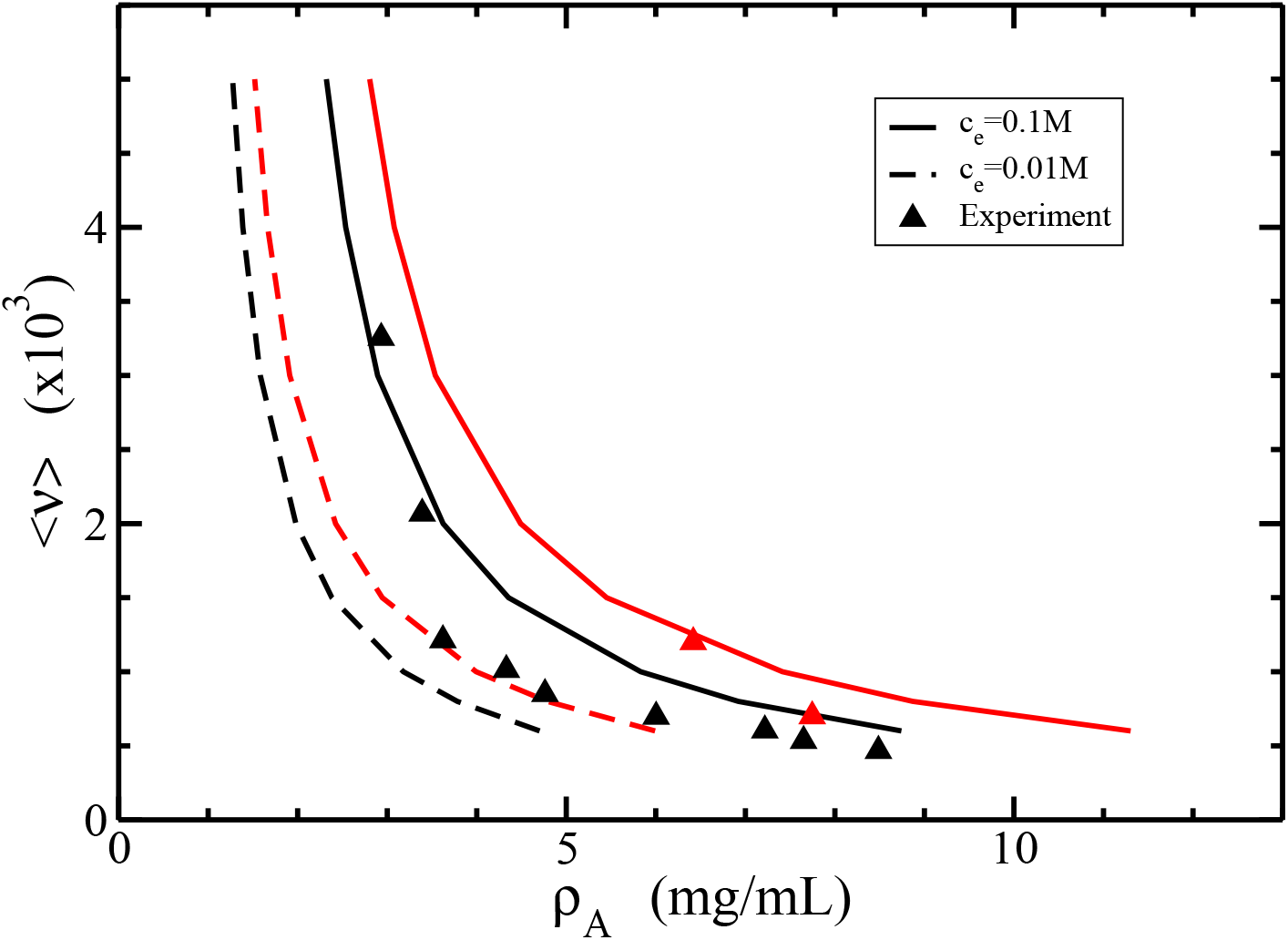
The isotropic-nematic phase diagram in term of coexisting densities of actin *ρ_A_* for different average length of filaments < *ν* > and two values of electrolyte concentrations *c_e_* = 0.01*M* (dashed lines), and *c_e_* = 0.1*M* (solid lines). The persistence length is *P* = 18*μm*, and the normalized standard deviation *σ* = 0.5. The experimental data (triangles) are retireved from Ref.[26]. The coexisting isotropic and nematic densities are plotted in black and red colors, respectively.

Clearly, the results for *c_e_* = 0.1*M* give the best fit against the experimental data. This behavior is due to a substantial decreases of the effective diameter *D_eff_* with increasing the electrolyte concentration *c_e_* (see Fig.10), as well as because the coexisting density *ρ_A_* is inversely proportional to the effective diameter (see Eq.(73)).

In Fig.12 we investigated the I-N phase diagram for a fixed average length < *ν* > in terms of the dependence of the coexisting densities of actin *ρ_A_* on the concentration of monovalent ions *c_e_*.

**Figure 12.**
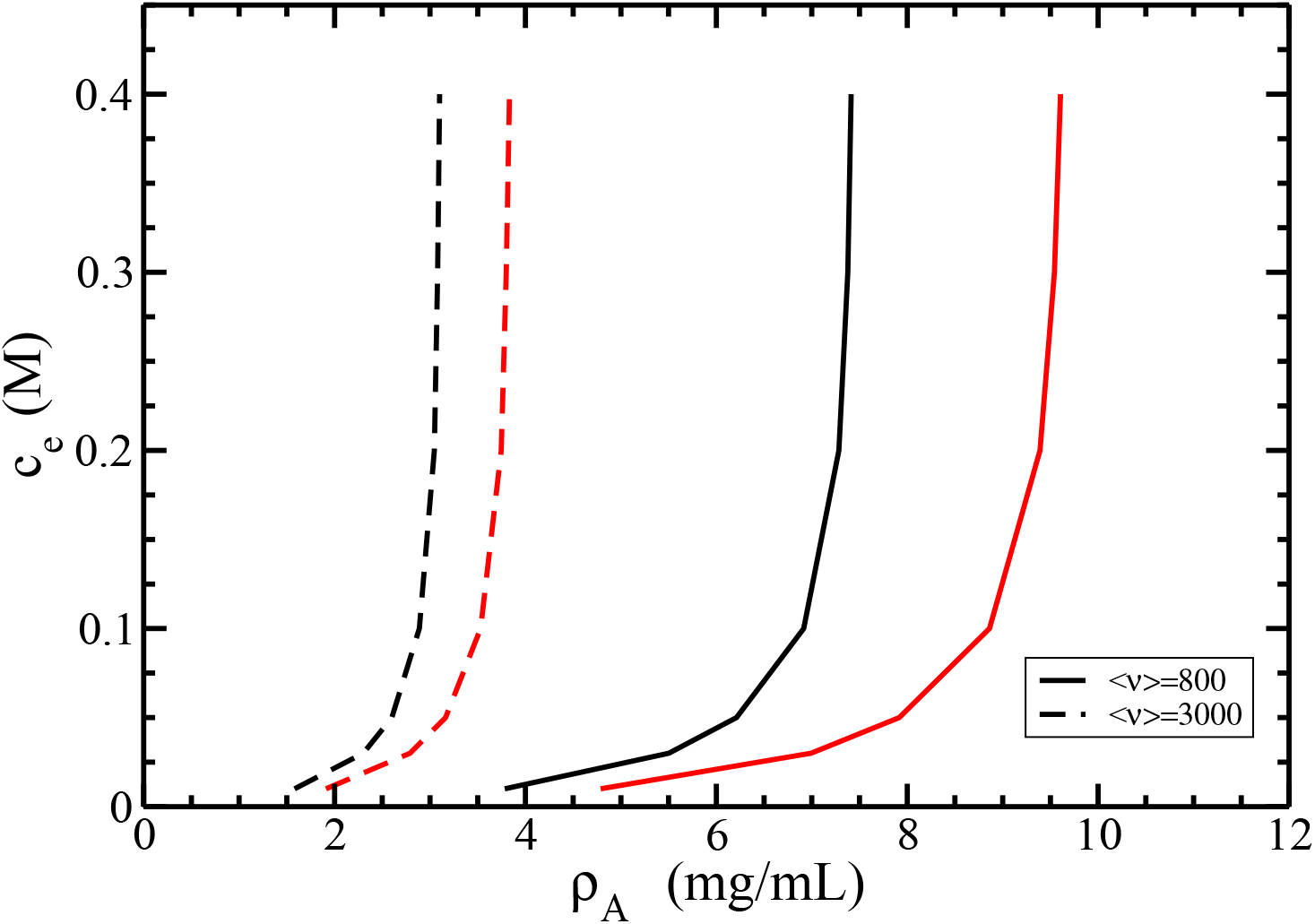
The isotropic-nematic phase diagram in term of coexisting densities of actin *ρ_A_* for different values of the monovalent ions concentrations *c_e_*. The average lengths of filaments are < *ν* >=800 (solid lines) and 3000 (dashed lines), the persistence length *P* = 18*μm*, and the normalized standard deviation *σ* = 0.5. The coexisting isotropic and nematic densities are plotted in black and red colors, respectively.

It is seen that the increase of the concentration *c_e_* leads to the increase of both isotropic and nematic coexisting densities *ρ_A_*. On the other hand, for fixed value of *c_e_*, the increase of average size < *ν* > decreases both coexisting densities *ρ_A_*. Certainly, it is easier for long rods or filaments to form a nematic order rather than for shorter ones. Alike, for longer filaments (< *ν* >= 3000), the I-N phase transition occurs at lower densities *ρ_A_* compared to short ones (< *ν* >= 800) with the same concentration *c_e_*.

Other quantity playing a key role in the I-N phase diagram is the twisting parameter. In some studies [32], the terms with exponential integral *E*_1_(*t*) in the formulas analogous to Eqs.(A14) and (64) were dropped off for simplicity, such as the electrostatic effects on the phase equilibrium can be described exclusively by the effective diameter *D_eff_* and the twisting parameter

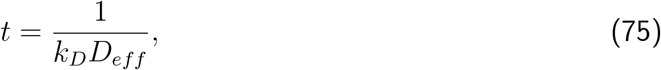

which is a measure of a twist of equally charged cylinders, which hinders the formation of the nematic order. On another hand, the increase of the effective diameter *D_eff_* promotes the nematic order. The competition between these two effects affects the I-N phase equilibrium in the electrolyte solution [32]. We plotted the dependence of the twisting parameter *t* on the concentration of monovalent ions *c_e_* in Fig.13.

**Figure 13.**
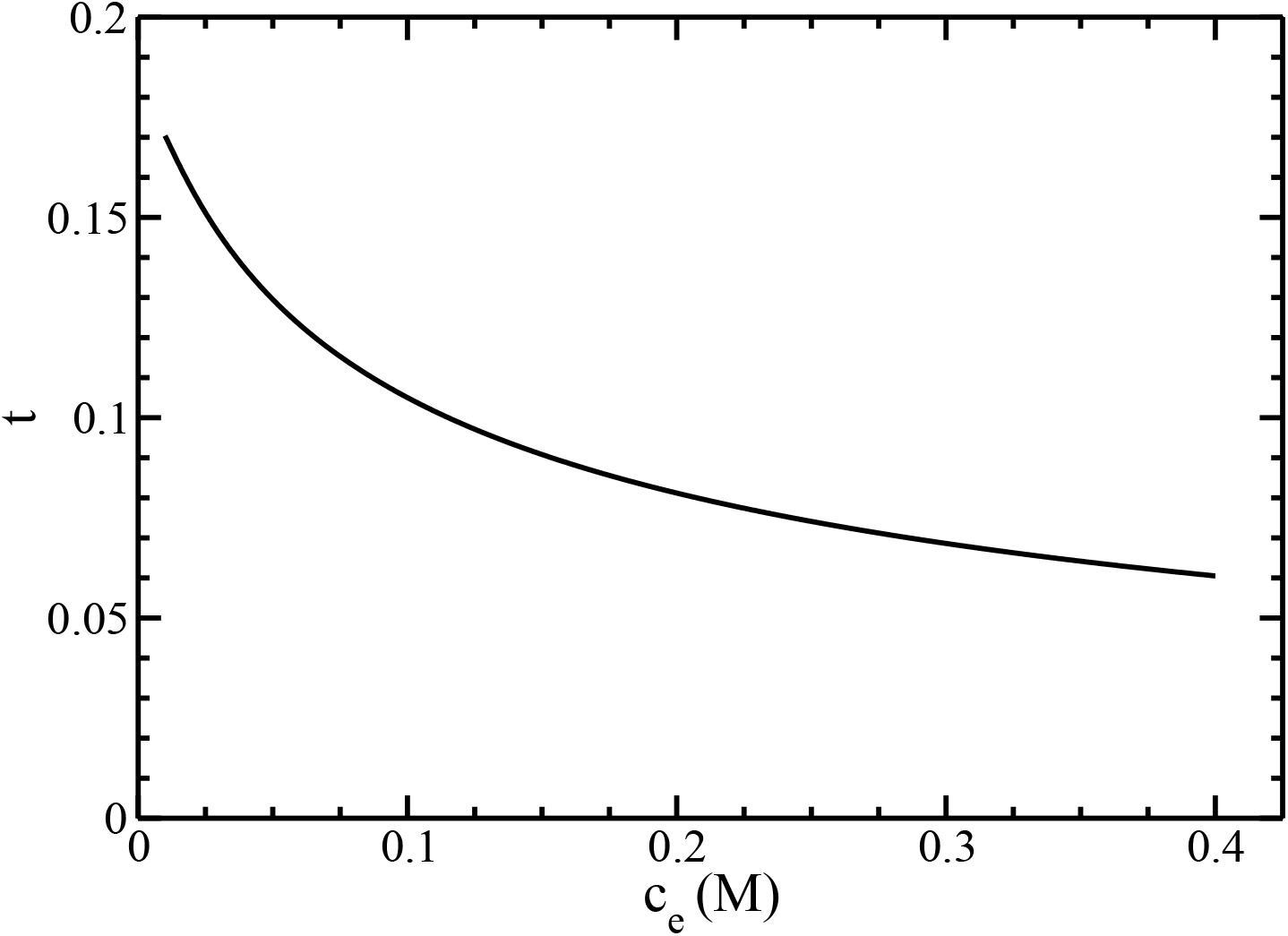
The twisting parameter *t* as a function of the monovalent ions ocnentration *c_e_*.

For actin filaments, the parameter *t* decreases with the increase of the concentration *c_e_*. We also rescale the I-N phase diagram in terms of coexisting densities of actin *ρ_A_* for different values of the twisting parameter *t* on Fig.14.

**Figure 14.**
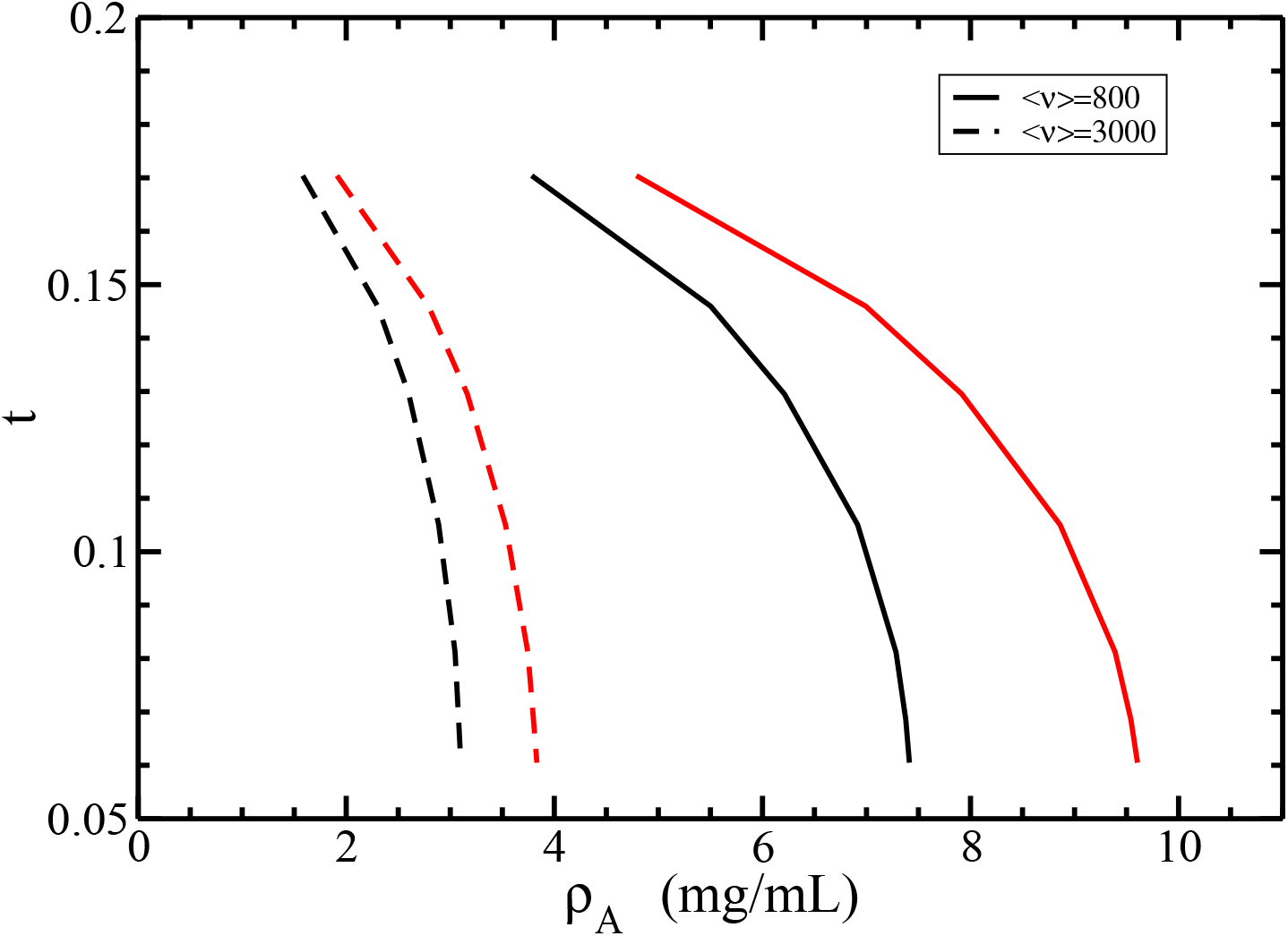
The isotropic-nematic phase diagram in term of coexisting densities of actin *ρ_A_* for different values of the twisting parameter *t*. The average lengths of filaments are < *ν* >=800 (solid lines) and 3000 (dashed lines), the persistence length *P* = 18*μm*, and the normalized standard deviation *σ* = 0.5. The coexisting isotropic and nematic densities are plotted in black and red colors, respectively.

It is seen that both isotropic and nematic coexisting densities *ρ_A_* decrease with increasing the twisting parameter *t*. Whereas, for a fixed value *t*, the increase of average size < *ν* > decreases both coexisting densities *ρ_A_*. Indeed, according to Figs.10 and 13, the increase of the twisting parameter *t* decreases the electrolyte concentration *c_e_* and, correspondingly, increases the effective diameter *D_eff_*. Furthermore, our results plotted in Fig.14 show that stabilizing effects on the nematic state of larger effective diameters *D_eff_* dominate destabilizing twisting effects with larger twisting parameters *t* [11].

## IV. DISCUSSION

In the present article we have developed a classical density functional theory to calculate the isotropic-nematic phase diagram for actin filaments in physiological electrolyte solutions. The approach is based on an extension of Onsager’s second order theory for monodisperse, charged rigid rods [10]. The key ingredients in this work are a unique definition of the orientational free energy *f_or_*, the *n_ν_* size and angular *η_ν_*(*θ*) distribution functions, which properly account for the polydispersity and semiflexibility of the actin filaments. Unlike many studies on polydisperse rods where the size distribution function *n_ν_* is an input parameter [17, 19], actin filaments self-aggregate to produce a size distribution function *n_ν_* that satisfies the equilibrium condition given by Eq.(27). Our results reveal that *n_ν_* has a form of Schulz’s distribution function, which is in accordance with experiment work done on actin filaments [25, 26]. To model an angular distribution function in nematic phase *η_ν_*(*θ*), we have generalized the trial function introduced in Ref.[17] for polydisperse filaments, which depends on two parameters α and γ. Additionally, we have accounted for the filament semiflexibility by extending the formula for orientational free energy *f_or_* introduced in Ref.[13] to the case of polydisperse system. Finally, we have calculated the isotropic-nematic phase diagrams for several values of the normalized standard deviation of Schult’s distribution *σ*, the persistence lengths of actin filaments *P*, and the concentrations of monovalent ions *c_e_*. We have compared the obtained results with the corresponding experimental data presented in Ref.[26]. We have found that the set of parameters *P* = 18*μm* and *c_e_* = 0.1*M* gives the best match against the experiment.

In principle, our method can be applied with suitable modifications to other associating particles or exchange colloids, which form charged rod-shaped aggregates. It can be also extended to study phase diagram in confined spaces, as well as under other electrolyte and filament conditions.

For instance, the orientational trial function used in this article is valid for weak polydispersity, i.e. when the normalized standard deviation *σ* is small.. However, the solution can also be obtained for highly polydisperse systems while the computational burden dramatically increases.

Additionally, in our study we followed the conditions of the experiment, where the average length of actin filaments < *ν* > was fixed both in isotropic and nematic phases [26]. As a result, the last two terms in the expression for free energy (43) did not contribute to the phase equilibrium. In principle, we can also apply our theory for polydisperse systems having different average lengths in the coexisting isotropic and nematic phases. In this case, the last two terms in the equation for free energy (43) can not be disregarded. Additionally, the approximation introduced in this work for the excess standard chemical potential 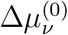 in Eq.(39) should be properly changed/modified if the length distribution of the polydisperse system does not follow the Schulz’s distribution function *n_ν_*.

On the other hand, we have used Brenner-Parsegian’s formula [27] to approximate the electrostatic part of inter-filament potential in Eq.(15). This approximation accounts for water solvent as a dielectric medium characterized by a constant dielectric permittivity. In principle, more accurate, distance-dependent dielectric medium can be used to capture the high water polarization near the filament surface [33].

Furthermore, actin filaments may form bundles and networks due to the attraction forces between filaments produced by linker proteins or divalent ions in eukaryotic cells. In our method we have accounted only repulsive interaction between filaments, i.e the hard core plus electrostatic repulsion in Eq.(14). The attraction between filaments can be included, for example, using square-well-type potentials [2, 34].

Our approach is also able to describe charged filaments in pathological conditions. For instance, pH changes and G-actin mutations have been accounted from molecular structure models for actin filaments [35].

Overall, more realistic models can be developed for filament-filament and filament-binding protein interactions in constrained spaces (encapsidated polyelectrolytes) [36, 37]. For instance, a variety of geometries such as spherical and cylindrical capsids can be used to model different cellular compartments such as dendrites (spines and filopodia), soma, axons, and terminals.

## Author Contributions

Conceptualization, MM; methodology, VV and MM; validation, VV; formal analysis, VV and MM; investigation, VV and MM; resources, MM; writing–original draft preparation, VV; writing–review and editing, MM; supervision, MM; project administration, MM; funding acquisition, MM. All authors have read and agreed to the published version of the manuscript.

## Institutional Review Board Statement

This study did not require ethical approval; all data are available in the public domain.

## Informed Consent Statement

Not applicable.

## Data Availability Statement

Some or all data, models, or code that support the findings of this study are available from the corresponding author upon reasonable request..

## Appendix A: Cluster integral *B*_*v*_1_*v*_2__(*ω*_1_,*ω*_2_) in Eq.(12)

The volume integration in Eq.(12) can be written as follow

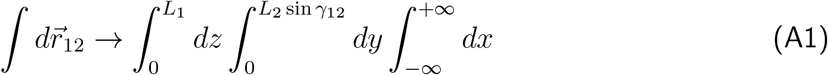

where the axis *x* is chosen along the direction of the closest distance between rods, the axis *z* along the direction of first rod, and the axis *y* in the perpendicular direction to *z* – *x* plane. *L*_1_, *L*_2_ are the lengths of two rods, and *γ*_12_ the angle between the axis of two rods. Substitution of Eq.(A1) into Eq.(12) yields

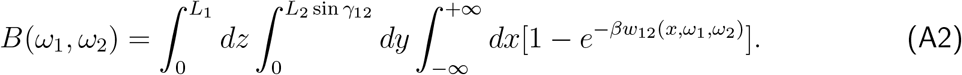

Integration in Eq.(A2) on the *y* – *z* plane leads to

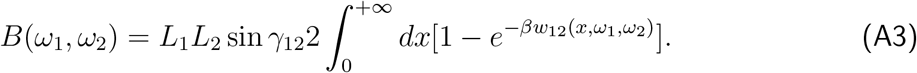

Substitution of Eq.(14) into Eq.(A3) yields

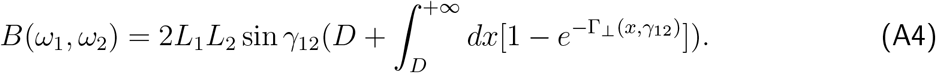

Substitution of Eq.(15) into the integral in the r.h.s of Eq.(A4) yields the following expression

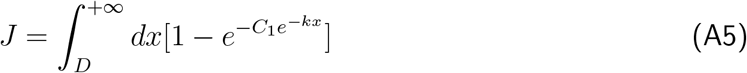

where

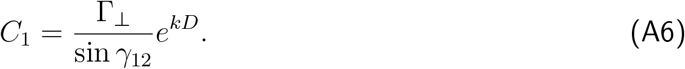

The analytic solution of Eq.(A5) is given by

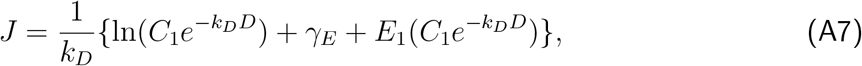

where *γ_E_* is the Euler’s constant

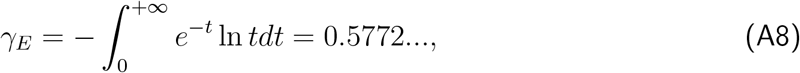

and *E*_1_(*x*) is the exponential integral

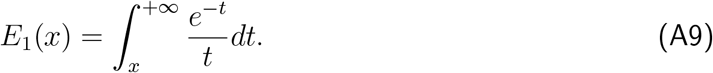

Next, we substitute Eqs.(A5),(A6),(A7) into Eq.(A4) to get

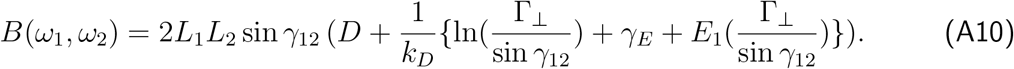

We introduce the effective diameter *D_eff_* to rewrite Eq.(A10) as follow

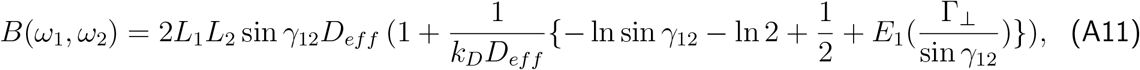

where

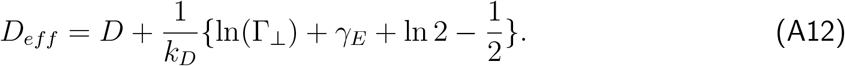

Finally, Eq.(A11) can be written as

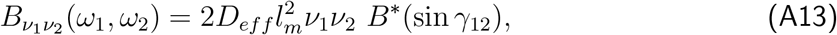

where

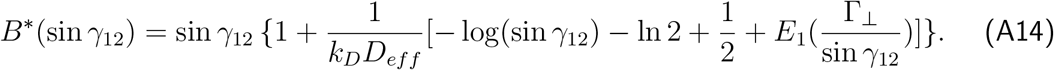

## Appendix B: Schulz’s distribution function in Eq.(40)

Here, we provide some properties of Schulz’s distribution function *n_ν_*. We define the average size in *k*-th power < *v^k^* > as follow

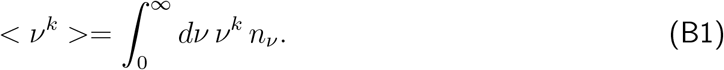

Substitution of Eq.(40) into Eq.(B1) gives

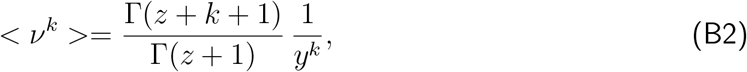

where Γ is the Gamma-function

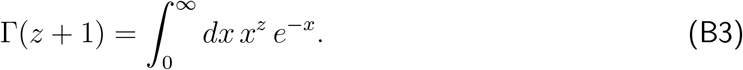

Eq.(B2) yields the following expressions for *k* = 0, 1, 2

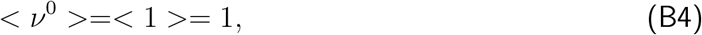

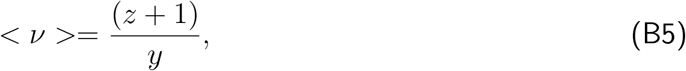

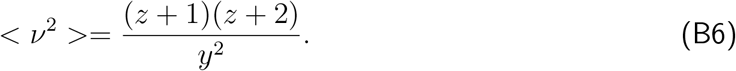

Substitution of Eqs.(B5) and (B6) in Eq.(5) generates the following relation between the normalized standard deviation *σ* and parameter *z*

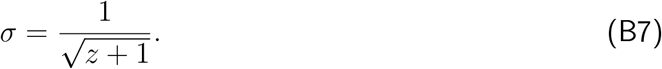

## Appendix C: Orientational energy *f_or_* in Eq.(49)

The orientational free energy *f_or_* in the nematic phase for monodisperse systems can be expressed as [13]

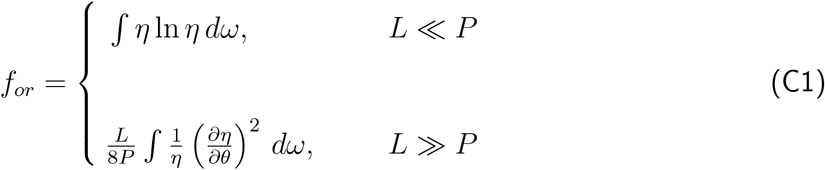

where *L* is the filament length, and *P* the persistence length. Khokhlov and Semenov introduced the correction terms to the r.h.s of Eq.(C1) to make this expression even more accurate [13–15]

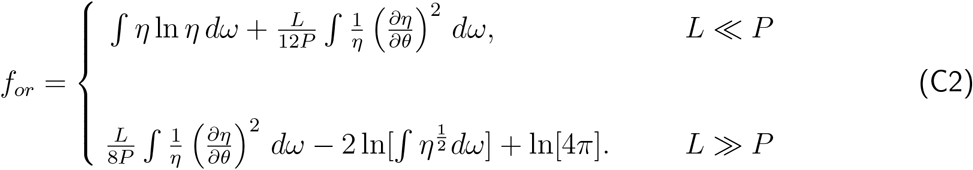

We generalize Eq.(C2) to the case of polydisperse system in the following form

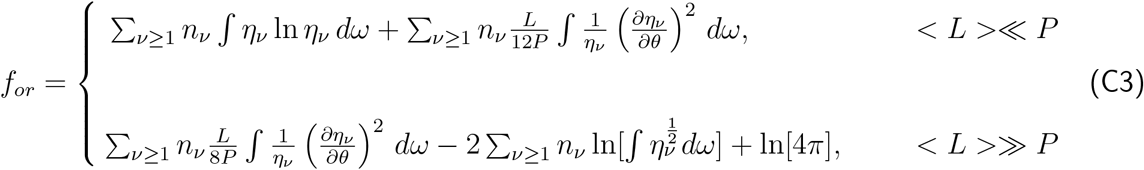

where < *L* >= *l_m_* < *v* >. Substitution of Eq.(41) into Eq.(C3) and the use of Eqs.(45) and (47) lead to

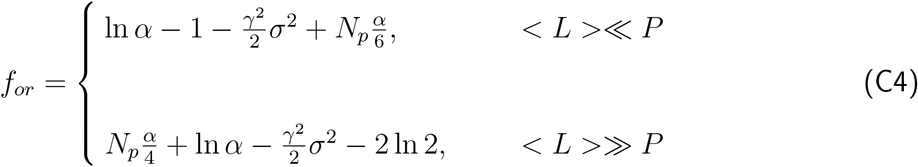

where *N_p_* is given by Eq.(48). We interpolate these two asymptotic values in Eq.(C4) to obtain

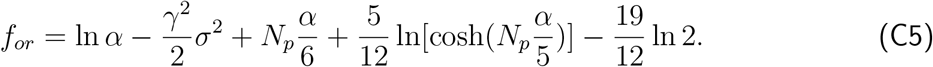

In monodisperse case, i.e. for *σ* = 0, Eqs.(48), (C4) and (C5) reduce to those provided in Ref.[13]. It was also suggested in Ref.[13] to substitute (*α* – 1)*N_p_* instead *αN_p_* in the monodisperse version of Eq.(C5) to have a reliable results for small *α* even when *N_p_* ~ 1.

## Appendix D: Interaction energy *f_int_* in Eqs. (50), (52), (53), and (54)

We substitute Eqs.(18),(19) into Eq.(50) to have

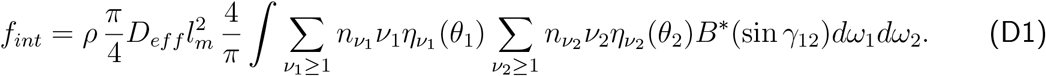

Eq.(D1) can be expressed as

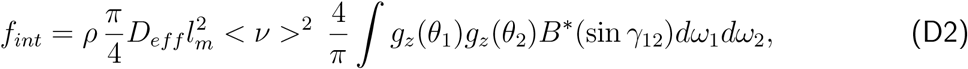

where

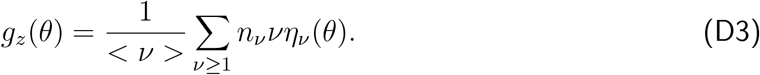

Additionally, we substitute Eq.(40) for the distribution functions *η_v_* and Eqs.(41) and (42) for *η_ν_*(*θ*) into Eq.(D3). It results

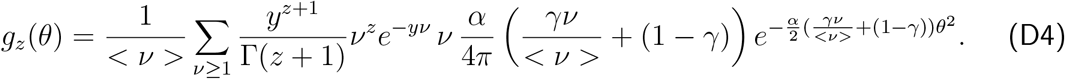

In Eq.(D4), the summation can be written in the form of integration as follow

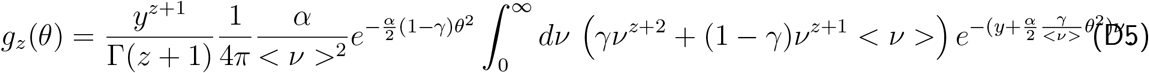

Using the definition of the Gamma-function given by Eq.(B3) Eq.(D5) becomes

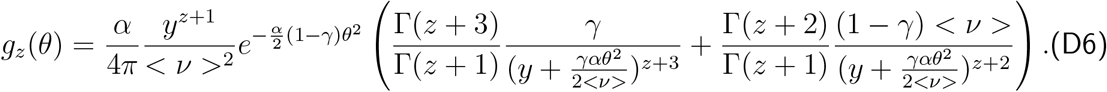

which can be rewritten as

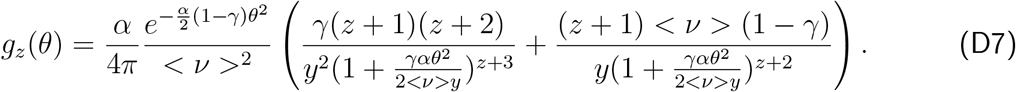

Substitution of the function *y* in Eq.(B5) into Eq.(D7) leads to

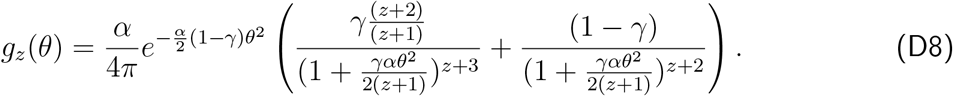

Finally, we transform the angular integrals in Eq.(D2). To this end we use the following

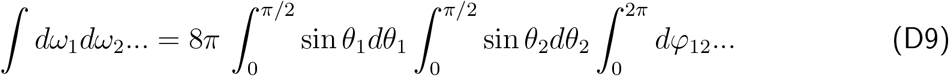

Substitution Eq.(D9) into Eq.(D2) yields

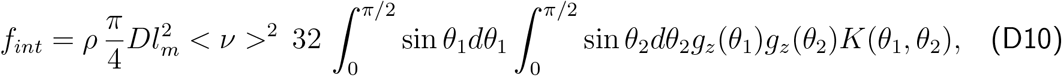

where

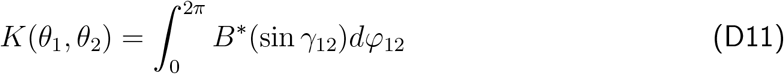

and

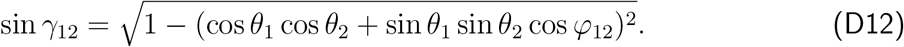

We note that the function *K*(*θ*_1_,*θ*_2_) is independent of the filaments concentration *ρ* and the average filaments length < *v* >. Thus, it can be numerically precalculated and used in subsequent calculations. In practice, we calculated *K* and stored the values in a matrix of size [200 × 200].

## Appendix E: Asymptotic expression for *f_mix_* in the monodisperse limit

We substitute the distribution function *n_v_*, given by Eq.(40), into Eq.(58). We get

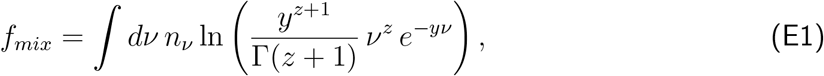

Substitution of Eqs.(B4),(40), and (B5) into Eq.(E1) yields

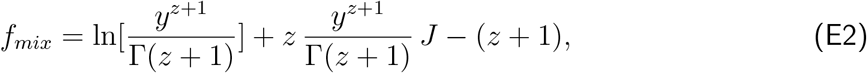

where

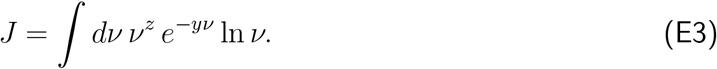

We obtain the analytic solution for *J* using the expression number 4.352 from the integral table provided in Ref.38. We have

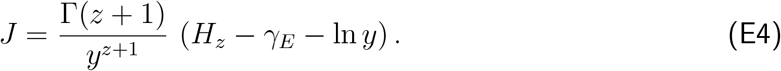

where *γ_E_* = 0.5772 is the Euler’s constant, Γ(*z* + 1) = *z*! the gamma-function, and 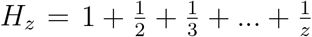 the *z*-th harmonic number. Substitution of Eq.(E4) into the second term in r.h.s of Eq.(E2) gives

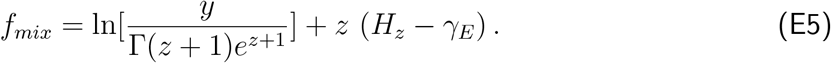

We use the following asymptotic formula for the harmonic number

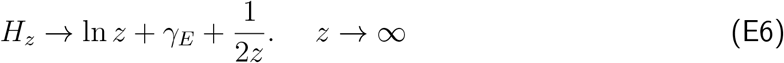

to calculate the free energy *f_mix_* in the monodisperse limit. We substitute Eq.(E6) into Eq.(E5), and we use Eqs.(B5) and (B7), and the Stirling’s formula 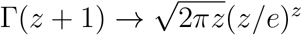 to finally obtain

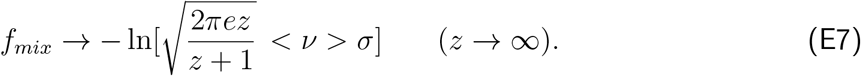

## References

[1] B Alberts, A Johnson, J Lewis, M Raff, K Roberts, and P Walter. Molecular Biology of the Cell, 6th Ed.,. Garland Science, New York, 2002.

[2] I Borukhov, R Bruinsma, W Gelbart, and A Liu. Structural polymorphism of the cytoskeleton: A model of linker-assisted filament aggregation. Proceedings of the National Academy of Sciences of the United States of America,, 102:3673–3678, 2005.

[3] A Suzuki, M Yamazaki, and T Ito. Osmoelastic coupling in biological structures: Formation of parallel bundles of actin filaments in a crystalline-like structure caused by osmotic stress. Biophysical Journal,, 28:6513–6518, 1989.

[4] A Suzuki, M Yamazaki, and T Ito. Polymorphism of f-actin assembly. 1. a quantitative phase diagram of f-actin. Biophysical Journal,, 35:5238–5244, 1996.

[5] G Wong, J Tang, A Lin, Y Li, P Janmey, and C Safinya. Hierarchical self-assembly of f-actin and cationic lipid complexes: stacked three-layer tubule networks. Science,, 288:2035–2039, 2000.

[6] J Xu, W H Schwarz, J A Kas, T P Stossel, P A Janmey, and T D Pollard. Mechanical properties of actin filament networks depend on preparation, polymerization conditions, and storage of actin monomers. Biophysical Journal,, 74:2731–2740, 1998.

[7] E D’Este, D Kamin, F Gottfert, A Al-Hady, and S W Hell. Sted nanoscopy reveals the ubiquity of subcortical cytoskeleton periodicity in living neurons. Cell Reports,, 10:1246–1251, 2015.

[8] S L Jones, F Korobova, and T Svitkina. Axon initial segment cytoskeleton comprises a multiprotein submembranous coat containing sparse actin filaments. Journal of Cell Biology,, 205:67–81, 2014.

[9] G Lukinavicius, L Reymond, E D’Este, A Masharina, F Gottfert, H Ta, A Guther, M Fournier, S Rizzo, H Waldmann, C Blaukopf, C Sommer, D W Gerlich, S W Arndt H D, Hell, and K Johnsson. Fluorogenic probes for live-cell imaging of the cytoskeleton. Nature Methods,, 11:731–733, 2014.

[10] L Onsager. The effects of shape on the interaction of colloidal particles. Annals of the New York Academy of Sciences,, 51:627–659, 1949.

[11] G Vroege and H Lekkerkerker. Phase transitions in lyotropic colloidal and polymer liquid crystals. Reports on Progress in Physics,, 55:1241–1309, 1992.

[12] A A. Stroobants, H N W. Lekkerkerker, and Th Odijk. Effect of electrostatic interaction on the liquid crystal phase transition in solutions of rodlike polyelectrolytes. Macromolecules,, 19:2232–2238, 1986.

[13] Th Odijk. Theory of lyotropic polymer liquid crystals. Macromolecules,, 19:2313–2329, 1986.

[14] A Khokhlov and A Semenov. Liquid-crystalline ordering in the solution of long persistence chains. Physica A,, 108:546–556, 1981.

[15] A Khokhlov and A Semenov. Liquid-crystalline ordering in a solution of partially flexible macro-molecules. Physica A,, 112:605–614, 1982.

[16] R Hentschke. Equation of state for persistent-flexible liquid-crystal polymers. comparison with poly(y-benzyl-l-glutamate) in dimethylformamide. Macromolecules,, 23:1192–1196, 1990.

[17] T Sluckin. Polydispersity in liquid crystal systems. Liquid Crystals,, 6:111–131, 1989.

[18] Z Y Chen. Effect of polydispersity on the isotropic-nematic phase transition of rigid rods. Physical Review E,, 50:2849–2855, 1994.

[19] A Speranza and P Sollich. Simpified onsager theory for isotropic-nematic phase equilibria of length polydisperse hard rods. Journal of Chemical Physics,, 117:5421–5436, 2002.

[20] H Wensink and G Vroege. Isotropic-nematic phase behavior of length-polydisperse hard rods. Journal of Chemical Physics,, 119:6868–6882, 2003.

[21] A Speranza and P Sollich. Isotropic-nematic phase equilibria in the onsager theory of hard rods with length polydispersity. Physical Review E,, 67:061702, 2003.

[22] W E McMullen, W M Gelbart, and A Ben-Shaul. Isotropic-nematic transition in micellized solutions. Journal of Chemical Physics,, 82:5616–5623, 1985.

[23] W E McMullen, W M Gelbart, and A Ben-Shaul. Translational and rotational contributions to the size of micelles in dilute soap solutions. Journal of Physical Chemistry,, 88:6649–6654, 1984.

[24] D Biron, E Moses, I Borukhov, and S A Safran. Inter-filament attractions narrow the length distribution of actin filaments. Europhysics Letters,, 73:464–470, 2006.

[25] J Drogemeier, H Hinsen, and W Eimer. Flexibility of f-actin in aqueous solution: A study on filaments of different average lengths. Macromolecules,, 27:87–95, 1994.

[26] A Suzuki, T Maeda, and T Ito. Formation of liquid crystalline phase of actin filament solutions and its dependence on filament length as studied by optical birefringence. Biophysical Journal,, 59:25–30, 1991.

[27] S Brenner and V Parsegian. A physical method for deriving the electrostatic interaction between rod-like polyions at all mutual angles. Biophysical Journal,, 14:327–334, 1974.

[28] R van Roij. The isotropic and nematic liquid crystal phase of colloidal rods. European Journal of Physics,, 26:S57–S67, 2005.

[29] B Zimm. Apparatus and methods for measurement and interpretation of the angular variation of light scattering; preliminary results on polystyrene solutions. Journal of Chemical Physics,, 16:1099–1116, 1948.

[30] P Janmey, J Peetermans, K Zaner, T Stossel, and T Tanaka. Structure and mobility of actin filaments as measured by quasielastic light scattering, viscometry, and electron microscopy. The Journal of Biological Chemistry,, 261:8357–8362, 1986.

[31] W H Press, S A Teulkolsky, V T Vetterling, and B P Flannery. Numerical Recipes in Fortran 77: The Art of Scientific Computing. Volume 1. Cambridge University Press 2nd Edition, Cambridge, New York, Melbourne, 1997.

[32] A Stroobants, N W Lekkerkerker, and Th Odijk. Effect of electrostatic interaction on the liquid crystal phase transition in solutions of rodlike polyelectrolytes. Macromolecules,, 19:2232–2238, 1986.

[33] V Warshavsky and M Marucho. Polar-solvation classical density-functional theory for electrolyte aqueous solutions near a wall. Physical Review E, 93:042607–19, 2016.

[34] A Grosberg and A Khokhlov. Statistical theory of polymeric lyotropic liquid crystals. Advances in Polymer Science,, 41:55–97, 1981.

[35] Christian Hunley, Md Mohsin, and Marcelo Marucho. Electrical impulse characterization along actin filaments in pathological conditions. Computer Physics Communications, 275:108317, 2022.

[36] Zhehui Jin and Jianzhong Wu. Density functional theory for encapsidated polyelectrolytes: A comparison with monte carlo simulation. The Journal of Chemical Physics, 137:(4), 044905, 2012.

[37] Ke Wang, Yang-Xin Yu, and Guang-Hua Gao. Density functional study on the structural and thermodynamic properties of aqueous dna-electrolyte solution in the framework of cell model. The Journal of Chemical Physics, 128:(18):185101, 2008.

[38] I S Gradshteyn and I M Ryzhik. Table of Integrals, Series, and Products, 5th Ed. Academic Press, Boston, MA, 1994.

